# Cargo-specific recruitment in clathrin and dynamin-independent endocytosis

**DOI:** 10.1101/2020.10.05.323295

**Authors:** Paulina Moreno-Layseca, Niklas Z. Jäntti, Rashmi Godbole, Christian Sommer, Guillaume Jacquemet, Hussein Al-Akhrass, Pauliina Kronqvist, Roosa E. Kallionpää, Leticia Oliveira-Ferrer, Pasquale Cervero, Stefan Linder, Martin Aepfelbacher, James Rae, Robert G. Parton, Andrea Disanza, Giorgio Scita, Satyajit Mayor, Matthias Selbach, Stefan Veltel, Johanna Ivaska

**Affiliations:** Turku Bioscience Centre, University of Turku and Åbo Akademi University, FI-20520 Turku, Finland; University Medical Center Hamburg-Eppendorf (UKE), 20251 Hamburg, Germany; National Centre for Biological Science (TIFR), Bellary Road, Bangalore, 560065, India; The university of Trans-Disciplinary Health Sciences and Technology (TDU), Yelahanka, Bangalore, 560064, India; Max-Delbrück-Centrum für Molekulare Medizin (MDC), 13092 Berlin, Germany; Faculty of Science and Engineering, Cell Biology, Åbo Akademi University, 20520 Turku, Finland; Institute of Biomedicine, Faculty of Medicine, University of Turku, FI-20520 Turku, Finland; Auria Biobank, Turku University Hospital and University of Turku, FI-20520 Turku, Finland; Institute for Molecular Bioscience, University of Queensland, Brisbane, QLD, 4072, Australia and Centre for Microscopy and Microanalysis, University of Queensland, Brisbane, QLD, 4072, Australia; IFOM, Fondazione Istituto FIRC di Oncologia Molecolare and University of Milan, Milan, 20139, Italy; Hochschule Bremen, City University of Applied Sciences, 28199 Bremen, Germany; Department of Biochemistry, University of Turku, FI- 20520, Turku, Finland

**Keywords:** Clathrin and dynamin-independent endocytosis, integrin, swiprosin-1, Rab21, breast cancer

## Abstract

Spatially controlled, cargo-specific endocytosis is essential for development, tissue homeostasis, and cancer invasion and is often hijacked by viral infections ^1^. Unlike clathrin-mediated endocytosis, which exploits cargo-specific adaptors for selective protein internalization, the clathrin and dynamin-independent endocytic pathway (CLIC-GEEC, CG-pathway) has until now been considered a bulk internalization route for the fluid phase, glycosylated membrane proteins and lipids ^2,3^. Although the core molecular players of CG endocytosis have been recently defined, no cargo-specific adaptors are known and evidence of selective protein uptake into the pathway is lacking ^3^. Here, we identify the first cargo-specific adaptor for CG-endocytosis and demonstrate its clinical relevance in breast cancer progression. By combining unbiased molecular characterization and super-resolution imaging, we identified the actin-binding protein swiprosin-1 (EFHD2) as a cargo-specific adaptor regulating integrin internalization via the CG-pathway. Swiprosin-1 couples active Rab21-associated integrins with key components of the CG-endocytic machinery, IRSp53 and actin. Swiprosin-1 is critical for integrin endocytosis, but not for other CG-cargo and supports integrin-dependent cancer cell migration and invasion, with clinically relevant implications for breast cancer. Our results demonstrate a previously unknown cargo selectivity for the CG-pathway and opens the possibility to discover more adaptors regulating it.

## Introduction

Endocytosis is a vital process of internalization of extracellular material and cell surface receptors that controls functions ranging from fluid-phase nutrient uptake, to spatially and temporally regulated traffic of adhesion and growth factor receptors and pathogen entry ^1^. The predominant view is that endocytosis specificity is achieved through cargo-specific adaptors, as described in detail for the best characterized internalization route clathrin-mediated endocytosis (CME) ^2,4^. This raises the possibility of yet unknown cargo adaptor proteins functioning as key gatekeepers for other endosomal routes. The CG-pathway internalizes a major fraction of the extracellular fluid phase, glycosylphosphotidylinositol (GPI)-anchored proteins and other cell surface receptors including nutrient transporters, ion channels and cell adhesion receptors ^5,6^ as well as pathogens such as bacteria and viruses ^5,7^. This non-specialized internalization of a variety of cargo occurs through high capacity tubulovesicular membrane uptake carriers called CLICs (clathrin-independent carriers) ^8,9^. CLICs are formed via the recruitment of Arf1, actin-binding BAR-domain protein IRSp53 and Arp2/3 to the membrane, followed by Cdc42 activation of IRSp53 and Arp2/3-mediated actin polymerization, presumptively resulting in the scission of CLICs and generation of CG endosomes ^10^. Although the core molecular players of CG endocytosis have been recently defined, no cargo-specific adaptors are known ^3^. The small GTPase Rab21 is unique in its ability to bind directly to integrin receptors to regulate endo/exosomal traffic, cytokinesis, chromosome integrity, endosomal signaling and anoikis ^11-14^. Rab21 interacts with integrins independently of its activation status (GDP/GTP); nevertheless, integrin endocytosis requires Rab21 activity ^11^ through currently unknown mechanisms. In addition, very few Rab21 interactors have been identified ^15^. Here, we identify swiprosin-1 (EFHD2) as a novel interactor of Rab21 and as the first cargo-specific adaptor for CG-endocytosis and demonstrate its clinical relevance as a prognostic marker for metastatic breast cancer.

## Results

We performed proteomic analyses by stable isotope labelling with amino acids in cell culture (SILAC) of Rab21 wild-type (WT), Rab21Q76L active mutant (CA-Rab21) or Rab21T31N inactive mutant (DN-Rab21) expressing cells to identify Rab21-interacting proteins ^16,17^. This mass spectrometry (MS) strategy identified the actin binding protein swiprosin-1 (EFHD2) as a putative active Rab21 interactor (Fig. 1A, Extended Data Fig. 1, Supplementary Table 1). Validation experiments with GFP pulldowns from cell lysate demonstrated endogenous swiprosin-1 binding preferably to WT and CA-Rab21, and not to the closely related Rab5 GTPase (Fig. 1B, Extended Data Fig. 2). In addition, purified recombinant GST-swiprosin-1 interacted directly with GTP-analog loaded recombinant Rab21 (Fig. 1C). The known Rab21-GTP specific interactor APPL1 ^8,19^ was included as a positive control. The interaction was further validated in cells. The proximity ligation assay (PLA) indicated endogenous swiprosin-1 and Rab21 interaction in intact cells (Fig. 1D), and bimolecular fluorescence complementation (BiFC) ^20^ revealed swiprosin-1 and Rab21 interaction at the cell-ECM interface (TIRF plane) in live cells (Fig. 1E, Extended Data Fig. 3A). The V1-Rab21/V2-swiprosin-1 BiFC interaction puncta were dynamic, detected at the cell periphery and in the cell center, and a similar signal was absent in Venus-expressing control cells (Supplementary Video 1). Together, these data identify swiprosin-1 as a novel active Rab21 interactor. Super-resolution structured illumination microscopy (SIM) images of the cell-ECM interface revealed overlap between swiprosin-1, Rab21 and β1-integrin in structures extending vertically into the cell (Fig. 1F-G, x-z projections). Swiprosin-1 co-localized significantly more with Rab21 than Rab5, Rab7 and Rab11 (Extended Data Fig. 2B), indicating a degree of specificity for Rab21 binding. Interestingly, Arf1 was detected as an active Rab21 interactor alongside swiprosin-1 (Fig. 1A, Extended Data Fig. 1) prompting us to test whether Rab21 and/or swiprosin-1 would associate with Arf1. Arf1 is a known regulator of CG-endocytosis (Fig. 1H), where cargo uptake is mediated by tubulovesicular membrane invaginations ^21^. Both GFP-Rab21 and GFP-swiprosin-1 co-immunoprecipitated endogenous Arf1 (Fig. 1I). In addition, another CG-endocytosis regulator, IRSp53 ^10^, was also detected in the co-immunoprecipitations (Fig. 1I). This indicates that the Rab21-swiprosin-1 complex is associated with the CG-endocytosis machinery. To investigate this more in detail, we imaged swiprosin-1 co-localization with known components of CME, caveolin-mediated endocytosis and CG-endocytosis using SIM. We observed swiprosin-1 co-localizing significantly more with Rab21, Arf1 and IRSp53 than clathrin, the clathrin adaptor AP2, caveolin or dynamin2 (Fig. 1J-K). These data are consistent with Rab21-mediated integrin endocytosis remaining unaffected by clathrin inhibition ^22^, the components of the clathrin and caveolin endocytic pathways not being enriched in the active Rab21 fractions analyzed by mass spectrometry (Fig. 1A, Extended Data Fig. 1) and the absence of dynamin I/II in bimolecular complementation affinity purified (BiCAP) ^20^ V1-Rab21 and V2-Swip1 complexes in cells (Fig. 1L; Extended Data Fig. 3B). Furthermore, Arf1 localization to swiprosin-1-positive structures was visible using another super-resolution microscopy technique: the localization-based method DNA-PAINT ^23^ (Extended Data Fig. 4) and Rab21/swiprosin-1-BiFC puncta were detected dynamically moving towards IRSp53-positive structures before disappearing from the TIRF plane in live cells (Fig. 1M, Supplementary Video 2). These data indicate the existence of a swiprosin-1/Rab21-GTP complex that links to the CG-endocytosis machinery.

**Fig. 1.**
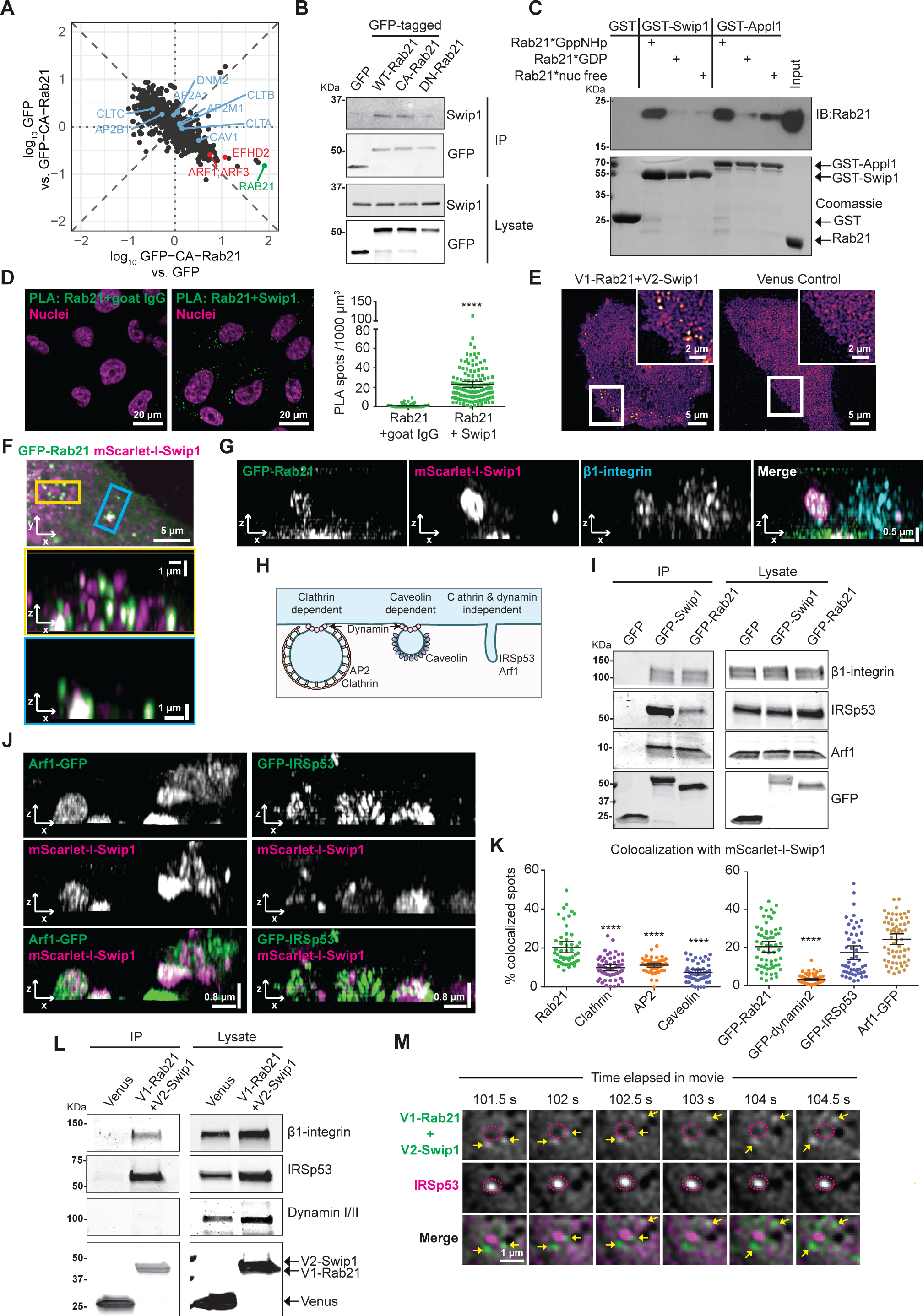
Swiprosin-1 (EFHD2, Swip1) links β1-integrin, the CG pathway and Rab21. **A**) SILAC proteomics analysis of GFP-TRAP pulldowns in MDA-MB-231 cells expressing GFP-CA-Rab21Q78L (constitutively active GTP-bound Rab21) vs GFP alone. The plot is representative of two independent experiments and every experiment consists of two independent affinity purifications. Each spot corresponds to one protein. Proteins in red are markedly enriched in the CA-Rab21 fraction (lower-right quadrant). Protein in blue are known endosomal proteins clathrin (CLTA, CLTB, CLTC), AP2 (AP2A1, AP2B1, AP2M1), caveolin (CAV1) and dynamin II (DNM2) and are not specifically enriched. **B**) Representative immunoblots of GFP-TRAP pulldowns from MDA-MB-231 cells transfected with the indicated constructs and probed for GFP and for endogenous swip1. GFP-DN-Rab21: Rab21T33N dominant negative. **C**) A coomassie-stained gel and immunoblot of GST pulldowns with the indicated GST-tagged proteins and recombinant Rab21 bound to a non-hydrolysable form of GTP (GppNHP; active Rab21), GDP or no nucleotide after EDTA treatment (nuc free). Recombinant GST-swip1 binds only to the active GTP-bound form of Rab21. The Rab21-effector GST-APPL1 is used as a positive control. **D**) Endogenous Rab21 and swip1 interact in MDA-MB-231 cells. A proximity ligation assay (PLA; green spots) with the indicated antibodies is quantified. Goat IgG was included as a negative control and nuclei were stained with DAPI. **E**) Representative TIRF microscopy BiFC images of live MDA-MB-231 cells expressing V1-Rab21 and V2-Swip1 or Venus alone as a control. **F**) MDA-MB-231 cells expressing mScarlet-I-Swip1 and GFP-Rab21 imaged using structured illumination microscopy (SIM). Blue and yellow squares highlight the regions of interest (ROI) in x-y that are magnified in x-z projections. **G**) SIM x-z projections of MDA-MB-231 cells expressing mScarlet-I-Swip1, GFP-Rab21 and immunostained for β1-integrin. **H**) Schematic of the major endocytic routes and their key components. **I**) Representative immunoblots of GFP-TRAP pulldowns from HEK293 cells transfected as indicated and blotted for GFP, endogenous IRSp53, Arf1 and β1-integrin. **J**) SIM x-z projections of MDA-MB-231 cells expressing mScarlet-I-Swip1 and GFP-IRSp53 or Arf1-GFP. **K**) MDA-MB-231 cells expressing mScarlet-I-Swip1 and immunostained for endocytic adaptor proteins or transfected as indicated, imaged with SIM and quantified for co-localization with mScarlet-I-Swip1. Each dot represents the co-localization ratio in one cell. **L**) Representative immunoblots of BiCAP pulldowns from MDA-MB-231 cells transiently transfected as indicated and blotted for GFP/Venus, endogenous IRSp53, dynamin I/II and β1-integrin (n = 2). The GFP antibody recognizes both V1-Rab21 and V2-Swip1 (see also Extended Data Fig. 3B). **M**) Representative TIRF microscopy BiFC images of live MDA-MB-231 cells expressing V1-Rab21+V2-Swip1 and mcherry-IRSp53. Yellow arrows point to V1-Rab21+V2-Swip1 puncta that travel towards an mcherry-IRSp53 puncta and then disappear from the TIRF plane. All immunoblots are representatives of three independent experiments unless stated otherwise. All plots show data from n = three experiments; mean ± 95% CI; **** P <0.0001, Mann Whitney test. Number of cells analyzed: (D) Rab21-IgG Goat, 121 and Rab21-Swip1, 126. (J) Rab21, 50; AP2, 50; caveolin, 50; clathrin, 55; GFP-Rab21, 63; GFP-dynamin2, 63; GFP-IRSp53, 63 and Arf1-GFP, 62.

CG-endocytosis is the major route for the uptake/internalization of versatile cargo: fluid phase endocytosis, the major histocompatibility complex I (MHCI) and cell surface receptors including integrins ^6^; whereas Rab21 has been primarily linked to integrin internalization ^11^, raising the possibility that the swiprosin-1/Rab21 complex could regulate specificity of integrin entry into the CG pathway. Rab21-integrin interaction requires the conserved KR-residues within the integrin α-subunit cytoplasmic tail ^11^ Mutation of these residues (KR1160/61AA) in the integrin α2-subunit ^11,14^ significantly reduced integrin colocalization with swiprosin-1-positive structures (Fig. 2A), indicating that a preserved integrin-Rab21 interaction is required for swiprosin-1 recruitment of the integrin receptor. We next investigated the requirement for swiprosin-1 in CG-endocytosis of integrins. Silencing of swiprosin-1 decreased, and overexpression of GFP-swiprosin-1 increased, the uptake of cell-surface labelled β1-integrin (antibody recognizes the active conformation of the receptor) in MDA-MB-231 cells. The effect on integrin endocytosis was apparent as early as the 5-minute time point and persisted for up to 30 minutes (Fig. 2B, Extended Data Fig. 5A). The expression of ectopic GFP-swiprosin-1 rescued β1-integrin endocytosis in swiprosin-1-silenced cells and increased the uptake in control siRNA transfected cells (Fig. 2C). Moreover, silencing of Arf1 or IRSp53, both essential members of the CG pathway, with two independent siRNAs, significantly decreased β1-integrin endocytosis in MDA-MB-231 cells (Extended Data Fig. 5B). Furthermore, integrin endocytosis was impaired in IRSp53-null mouse embryonic fibroblasts (IRSp53/ MEF; Extended Data Fig. 5C). This was specifically due to reduced integrin uptake and not due to lower integrin levels, since total integrin levels were unaffected by loss of IRSp53 or swiprosin-1 (Extended Data Fig. 5C-D). Swiprosin-1-induced integrin endocytosis was dependent on the CG pathway machinery, since depletion of IRSp53 abolished the ability of GFP-swiprosin-1 overexpression to induce β1-integrin endocytosis (Fig. 2D). In addition, the initial uptake of integrins overlapped with other CG cargos. Immediately after endocytosis (2 and 5 minutes), plasma membrane labelled β1-integrin co-localized with 10 kDa dextran, a fluid phase cargo for the CG pathway, as efficiently as the co-localization of dextran labelled with two different dyes (Fig. 2E). Taken together, these data indicate that swiprosin-1 induces integrin endocytosis via the CG pathway in these cells.

**Fig. 2.**
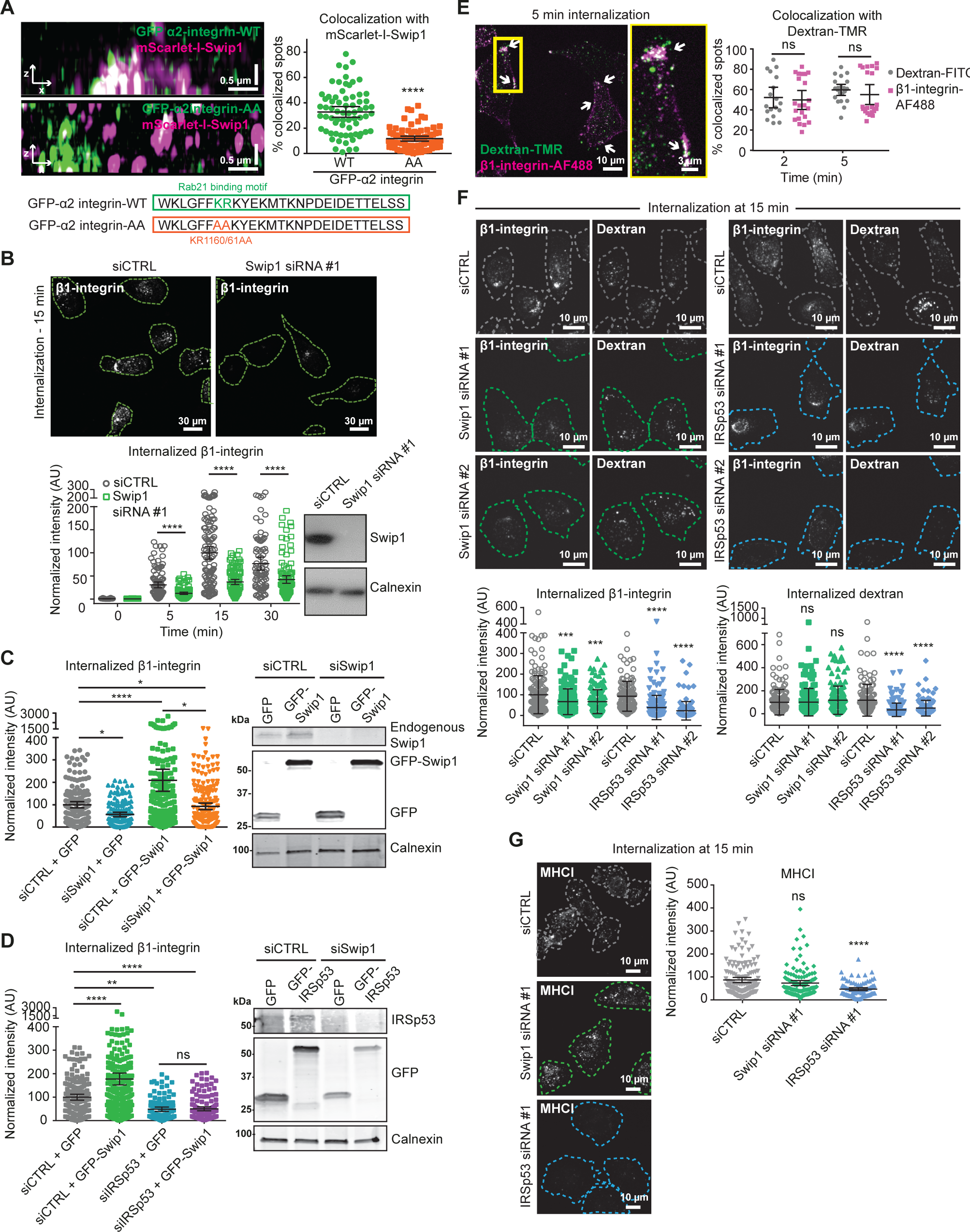
Swip1 is a cargo-selective adaptor for the CG pathway. **A**)SIM x-z projections of MDA-MB-231 cells expressing mScarlet-I-Swip1 and GFP-tagged α2 integrin-WT or mutant α2 integrin-AA (deficient in Rab21 binding; the integrin cytoplasmic sequence and mutated residues are depicted). Co-localization between GFP and mScarlet-I-Swip1 was quantified. **B**) Representative micrographs of β1-integrin internalization in control- and swip1-silenced MDA-MB-231 cells (siRNA 1) and quantification of integrin internalization at 5, 15 and 30 minutes. Representative immunoblot to validate swip1 silencing is shown. **C-D**) Quantification of β1-integrin internalization at 15 min in (C) control- or swip1-silenced MDA-MB-231 cells (siRNA#1) expressing GFP or GFP-Swip1 or in (D) control- or IRSp53-silenced MDA-MB-231 cells (siRNA#2) expressing GFP or GFP-Swip1. Representative immunoblots of cell lysates probed as indicated. **E**) Double uptake of fluorescently labelled 10 kDa dextran-TMR (tetramethylrhodamine) with either dextran-FITC or Alexa Fluor 488-conjugated anti-β1-integrin antibody (12G10) in MDA-MB-231 cells for the indicated times. **F**) Representative micrographs and quantification of dextran-TMR or anti-β1-integrin antibody (conjugated to Alexa Fluor 488) internalization in control-, swip1- (siRNA1 or 2) or IRSp53- (siRNA#1 or #2) silenced MDA-MB-231 at 15 min. **G**) Representative micrographs and quantification of Major Histocompatibility Complex I (MHCI) internalization in control-, swip1- (siRNA1) or IRSp53- (siRNA1) silenced MDA-MB-231 at 15 min. All immunoblots are representatives of three independent experiments and calnexin is included as a loading control. (B-G) Data are mean ± 95% CI; n = three experiments; *P <0.05, ***P <0.005, ****P <0.0001, ns: not significant, Mann Whitney test. Number of cells analyzed: (A) α2 integrin-WT, 70 and mutant α2 integrin-AA, 76. (B) siCTRL, 86, 101, 110 and 78; Swip1 siRNA#1, 104, 98, 102 and 106; from left to right. (C) siCTRL+GFP, 153; siCTRL+GFP-Swip1, 186; siSwip1+GFP, 136 and siSwip1+GFP-Swip1, 196. (D) siCTRL+GFP, 157; siCTRL+GFP-Swip1, 289; siIRSp53+GFP, 108 and siIRSp53+GFP-Swip1, 196. (E) Dextran-FITC, 18 and 22 and β1-integrin-A488, 21 and 21. (F) siCTRL, 160; Swip1 siRNA#1, 157; Swip1 siRNA#2, 160; siCTRL, 121; IRSp53 siRNA#1, 177 and IRSp53 siRNA#2, 121. (G) siCTRL, 152; Swip1 siRNA#1, 125 and IRSp53 siRNA#1, 92.

Next, we investigated whether swiprosin-1 regulates the endocytosis of other CG cargos. Swiprosin-1 silencing had no effect on the uptake of 10 kDa dextran or MHCI (Fig. 2F-G). In contrast, silencing of IRSp53 significantly impaired endocytosis of both cargos (Fig. 2F), validating the approach. In addition, swiprosin-1 depletion did not affect the endocytosis of transferrin or EGFR (Extended Data Fig. 5E-F), which are internalized through CME or dynamin-dependent non-clathrin endocytosis ^24,25^. Taken together these data imply that swiprosin-1 specifically controls integrin cargo recruitment towards CG-endocytosis rather than affecting the overall activity of this pathway. These data explain the long-standing conundrum of the requirement for active Rab21 in integrin endocytosis ^11^. Even though Rab21 binds integrins in GDP and GTP-bound states, swiprosin-1 interacts specifically with Rab21-GTP, coupling active Rab21 and integrin cargo to CG-endocytosis. A common emerging theme among non-clathrin endocytosis is a reliance on the actin cytoskeleton ^26^. Intrigued by the established actin-binding activity of swiprosin-1 ^27^, we investigated whether this function was important for swiprosin-1-mediated integrin CG-endocytosis. Deleting the first EF-hand domain (EF1) of swiprosin-1, rendered swiprosin-1 unable to bind actin, concordant with a previous report ^27^, and abolished its ability to induce integrin endocytosis (Fig. 3A-C). In contrast, EF2 deletion had no significant effect on swiprosin-1-induced integrin uptake. This indicates that swiprosin-1 binding to actin is necessary for its ability to augment integrin endocytosis. SIM imaging revealed F-actin overlap with swiprosin-1 and β1-integrin close to the cell-ECM interface (Fig. 3D), indicating that swiprosin-1 and the actin cytoskeleton are in close proximity during integrin endocytosis. In addition to swiprosin-1 localization at cell-ECM proximal structures, we observed swiprosin-1 deeper inside the cell, where it overlapped with Rab21-positive endosome-like vesicles (Fig. 3E) and F-actin in discrete puncta around Rab21 vesicles (Fig. 3E yellow arrows, Supplementary Video 3). Similar localization was visualized using GBP-APEX (GFP-binding protein soybean ascorbate peroxidase) labelled GFP-swiprosin-1 imaged with electron microscopy (Fig. 3F) ^28^. Swiprosin-1 localized to filaments close to the plasma membrane and in vicinity of endosomes (Fig. 3E pink arrow and 3F blue arrows). Swiprosin-1 localization with actin on Rab21 endosomes led us to investigate whether swiprosin-1 regulates endosome movement. Silencing of swiprosin-1 significantly reduced the speed of GFP-Rab21 vesicle movement and restricted their subcellular distribution to the cell periphery (Fig. 3G, Supplementary Videos 4-5). The motility of Rab21 vesicles was actin dependent, since the actin inhibitor cytochalasin D reduced vesicle speed (Fig. 3H), consistent with previous observations ^11^. Taken together, these data highlight a role for actin in both swiprosin-1/Rab21-dependent integrin CG-endocytosis and Rab21-mediated integrin endosomal traffic within the cell (Fig. 3I).

**Fig. 3.**
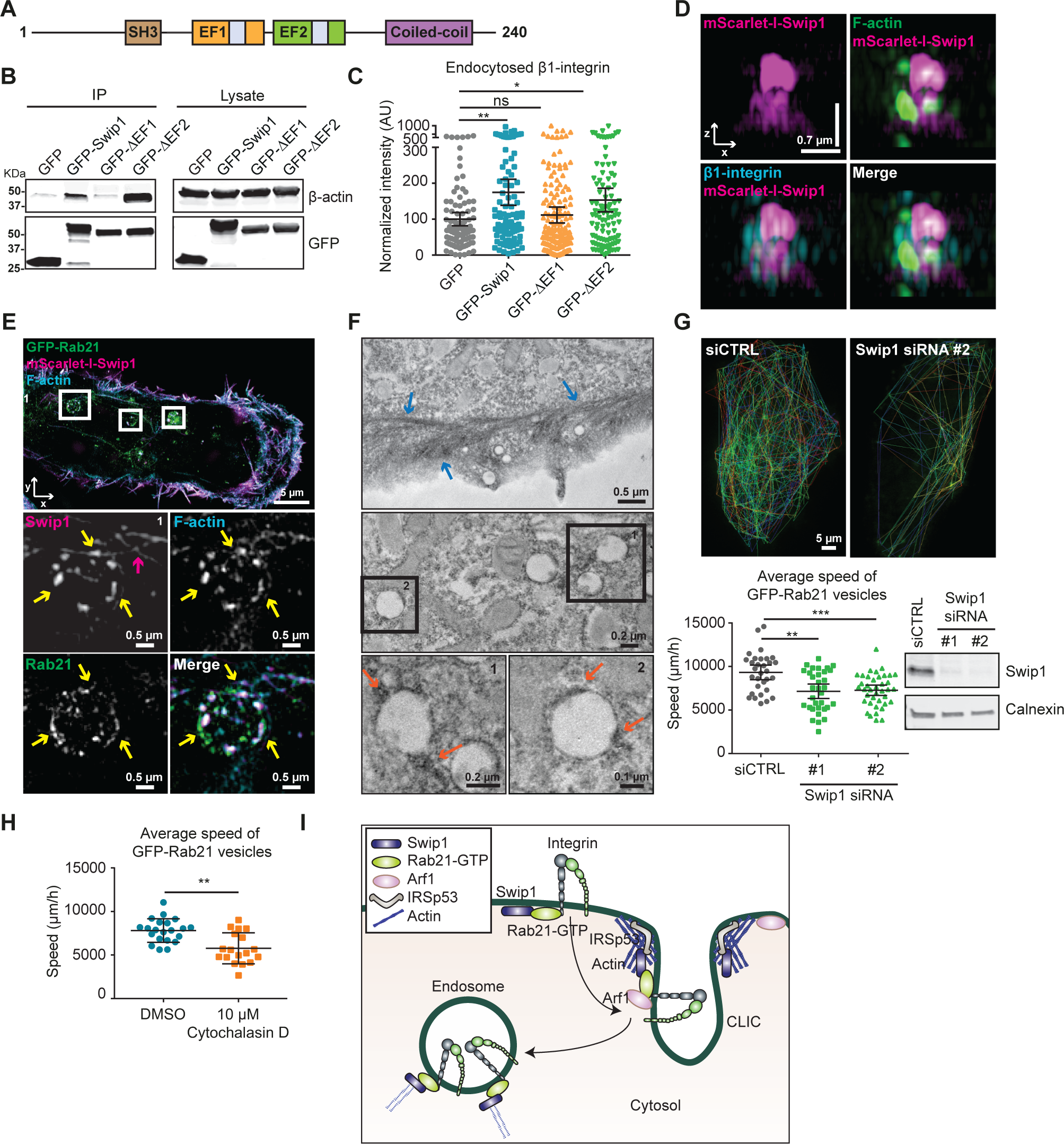
Swip1-actin binding regulates integrin traffic. **A**) Swip1 domains. EF: EF-hand domain containing a calcium-binding site (shown in gray). **B**) Representative GFP-pulldowns in HEK293 cells expressing GFP, GFP-swip1, or the truncated versions of swip1 and blotted for β-actin and GFP. **C**) Quantification of β1-integrin uptake at 15 min in MDA-MB-231 cells expressing either GFP, GFP-swip1 or the truncated versions of swip1 and normalized relative to GFP cells. **D**) SIM x-z projections of MDA-MB-231 cells expressing mScarlet-I-Swip1, immunostained for β1-integrin and labelled with phalloidin (F-actin). **E**) Representative SIM x-y image of MDA-MB-231 cells expressing mScarlet-I-Swip1, GFP-Rab21 and immunostained with phalloidin. White squares highlight ROIs. ROI1 is magnified. Yellow arrows point to swip1 overlap with actin on Rab21-containing vesicles. Pink arrow indicates actin filaments in close proximity to the vesicle. **F**) Electron microscopy images of GFP-Swip1 visualized using GBP-APEX. Squares highlight ROIs, which are magnified. Arrows point to swip1-APEX-positive patches adjacent to filament-like actin structures (blue arrows) or vesicles (orange arrows). **G**) Average speed of Rab21 vesicles, per cell, close to the TIRF plane over 2 min. Representative tracks of Rab21 vesicles in a control or a swip1-silenced cell and an immunoblot validating swip1 silencing are shown. Calnexin is included as a loading control. **H**) Average speed of Rab21 vesicle movement per cell upon treatment with cytochalasin D. **I**) Swip1 directs integrins to CG endocytosis. All representative immunoblots are from 3 independent experiments. Data are mean ± 95% CI; n = three experiments; *P<0.05, **P<0.005, ***P<0.0005, Mann Whitney test. Number of cells analyzed: (C) GFP, 92; GFP-Swip1, 94; GFP-1, 161 and GFP-2, 101. (G) siCTRL, 31; siSwip1 siRNA#1, 33; siSwip1 siRNA#2, 39. (H) DMSO, 21 and Cytochalasin D, 18.

Integrin endocytosis and intracellular transport are crucial for integrin turnover, cell migration and invasion ^15,29^. Concordant with this notion we found that vinculin-containing focal adhesions accumulated in swiprosin-1-silenced cells on collagen I, fibronectin and laminin-1-coated surfaces (Fig. 4A). This phenotype was also present in cells silenced for Arf1 or IRSp53 (Extended Data Fig. 6) indicating that integrin endocytosis through the CG pathway may function to regulate adhesion dynamics and cell motility. Using live-cell imaging of paxillin-positive focal adhesions, we observed significantly slower adhesion assembly and disassembly rates in swiprosin-1-silenced cells compared to control cells (Fig. 4B). This correlated with significantly impaired cell migration in a 2D scratch wound assay (Fig. 4C), in line with previously reported migration defects of IRSp53-null fibroblasts ^30^, and defective cell invasion through a 3D collagen matrix (Fig. 4D). These data indicate that swiprosin-1 stimulates integrin adhesion dynamics and the migration and invasion of breast cancer cells. This may have clinically relevant implications in breast cancer. qPCR analyses of swiprosin-1 mRNA levels from 192 breast cancer specimens revealed that swiprosin-1 expression was significantly higher in the highest grade tumors and the most metastatic breast cancer subtypes: HER2-positive and triple negative (TNBC) (Extended Data Fig. 7). These findings were further validated with immunohistochemistry of HER2+ and TNBC tissue microarrays using a swiprosin-1 specific antibody (Extended Data Fig. 8A). Swiprosin-1 was highly expressed in a high proportion of both breast tumour subtypes (65-75%), and in the case of TNBC, high swiprosin-1 staining in our cohort was associated with poor clinical outcome (Fig. 4E). We also analysed the prognostic value of high membranal signal for swiprosin-1 and found a more pronounced correlation with poor clinical outcome (Extended Data Fig. 8B-D). Moreover, we observed that the TNBC patients with high membranal swiprosin-1 had significantly more lymph node metastasis than that of medium-low membranal swiprosin-1 expressing patients (Fisher’s exact test, p=0.037). According to the Cox proportional hazards model, high swiprosin-1 membranal staining was associated with poor prognosis after adjustment with Ki67, tumor size, lymph node metastasis status, or grade (Extended Data Fig. 8). These data, demonstrating that elevated swiprosin-1 levels strongly and independently correlate with breast cancer metastasis and reduced survival, advocate swiprosin-1 as an original, independent prognostic marker and a potential drug target.

**Fig. 4.**
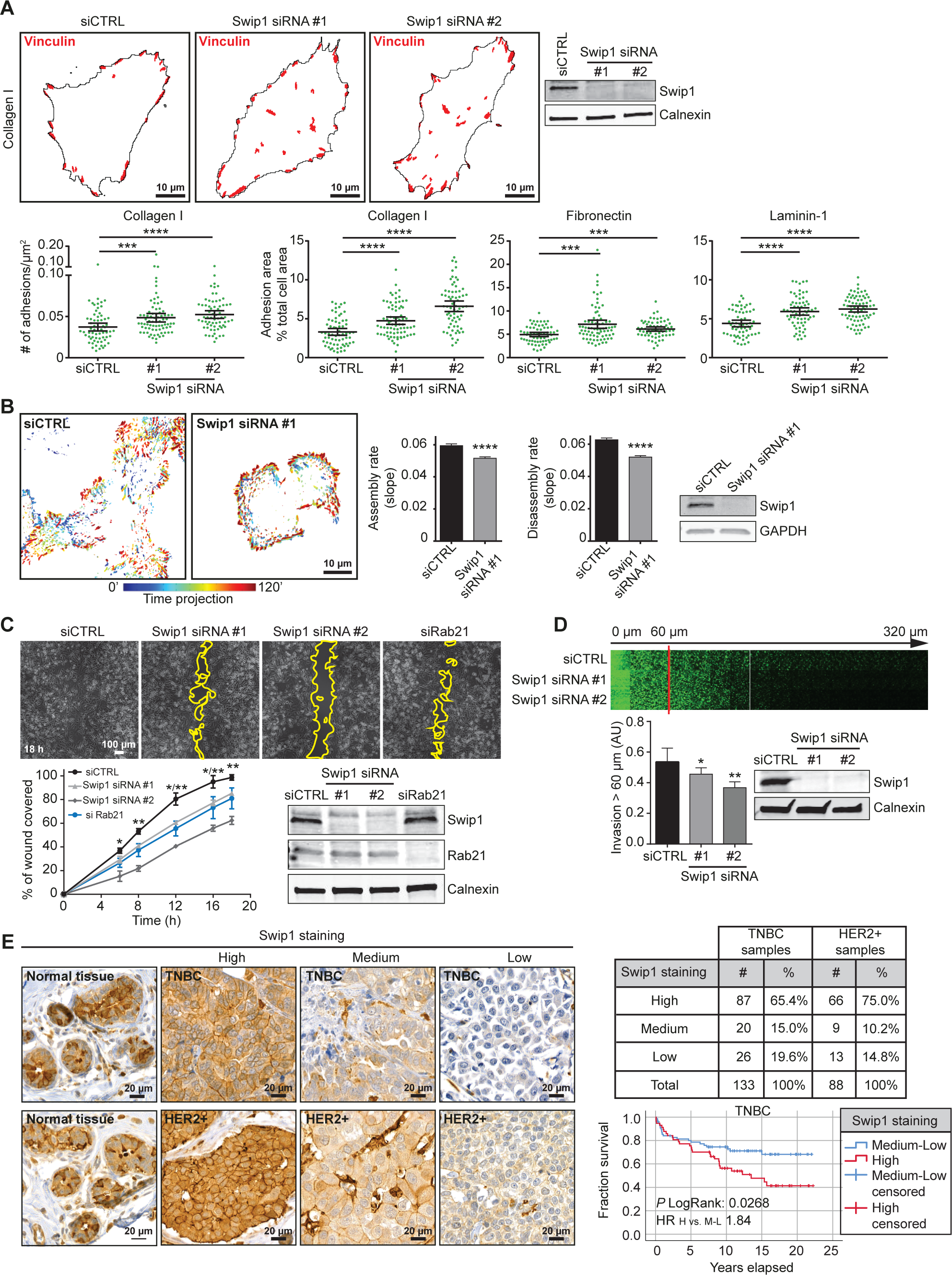
Swip1 regulates adhesion dynamics, cell motility and breast cancer progression. **A**) Representative masks of vinculin-containing focal adhesions in control- or swip1-silenced MDA-MB-231 cells on collagen I. Analysis of adhesion number and total adhesion area per cell on different ECM components. Representative immunoblot validating swip1 silencing is shown and calnexin is included as a loading control. **B**) Visualization of GFP-paxillin dynamics in control- and swip1-silenced cells. Color scale represents focal adhesion localization over time (120 min). Focal adhesion assembly and disassembly rates are quantified. Representative immunoblot validating swip1 silencing is shown, calnexin is included as a loading control. **C**) Migration of control-, swip1- or Rab21-silenced MDA-MB-231 cells. Representative phase contrast pictures of the scratch wound after 18 h and quantification of the wound area coverage over time (mean ± SEM, n = 3 experiments) are shown. Representative immunoblots validating swip1 and Rab21 silencing are shown, calnexin is included as a loading control. **D**) Control- and swip1-silenced MDA-MB-231 cells were seeded into an inverted invasion assay for 72 h. The relative invasion through fibrillar collagen I over 60µm was quantified. Representative immunoblot validating swip1 silencing is shown, calnexin is included as a loading control. **E**) images of swip1 staining in samples of HER2+ and TNBC breast cancer tissue microarrays and quantification of the percentage of tumors with high, medium or low swip1 are shown. Kaplan Meier plot shows overall survival of 133 TNBC patients with high (red) or medium and low (blue) staining of swip1. Hazard ratio (HR) was 1.84 (95 % CI 1.05 to 3.23). All representative immunoblots are from 3 independent experiments. Data are mean ± 5 % CI (A) or mean ±SEM (B, D); n = three experiments; *P<0.05, **P<0.005, ***P<0.0005, Mann Whitney test. Number of analyzed cells: (A) collagen I, 65, 67 76; fibronectin, 65, 75 62; laminin-1, 62, 66 73 for siCTRL, siSwip1 siRNA#1 and siRNA#2, respectively. (B) 18 cell per condition; number of FAs analyzed: siCTRL, 861 764; and siSwip1, 1014 1024 for assembly and disassembly, respectively.

## Discussion

Swiprosin-1 has not been previously associated with endocytosis and evidence for specific cargo adaptors or selective uptake in the CG-endocytosis pathway has been lacking. Our findings place swiprosin-1 as the first validated cargo-specific adaptor for the CG pathway. Swiprosin-1 directs active Rab21 and β1-integrins to the CG pathway and is necessary for active integrin endocytosis through this route, while fully dispensable for the uptake of other CG cargos. This demonstrates an important and unexpected feature of this pathway: cargo-specific adaptor-based recruitment of receptors. It also highlights the possibility of unprecedented cargo selectivity functions for the CG pathway in a manner similar to that described for numerous other endocytosed proteins following different endocytosis routes. Moreover, our findings explain the previously reported link between the CG pathway components, IRSp53 and Arf1, and adhesion phenotypes: IRSp53 contributes to cell migration and invasion through multiple mechanisms, including filopodia generation ^31,32^, and curvature sensing with WAVE at the neck of membrane invaginations ^33^. Arf1 depletion results in defective adhesion and migration in MDA-MB-231 cells ^34-36^, and our finding of Arf1 involvement in integrin endocytosis sheds light on the mechanism behind these observations. Interestingly, knockout of swiprosin in Drosophila results in a mild adhesion defect similar to the deletion of Rho-GAP-domain-containing protein GRAF1, another regulator of CLIC formation, suggesting an in vivo relevant role for swiprosin-1 in regulating cell-ECM interactions ^37,38^. Our data explains why Rab21 activation is required for integrin endocytosis and defines the thus far unknown Rab21-regulated endocytosis route. How Rab21 is activated, possibly by integrin-mediated adhesion, is an area that requires further investigation. Recently, a proximity-biotinylation screen revealed that Rab21 regulates sorting of a subset of clathrin-independent cargos ^39^. However, the involvement of swiprosin-1 in later endosomal sorting or retromer-mediated recycling ^39^ remains to be investigated. Here, we showed that swiprosin-1 acts as a cargo adaptor bridging the CG endocytic machinery to Rab21-bound integrin cargo, and couples Rab21 endosomes, and their motility in cells, to the actin cytoskeleton. This dual functionality of swiprosin-1 regulates cell adhesion turnover, migration and invasion. Perhaps unsurprisingly, given the importance of these events in cancer progression, swiprosin-1 expression has clinically relevant implications for TNBC. Thus, swiprosin-1 and the mechanism of its interaction with Rab21 offer potentially exciting therapeutic targets for metastatic breast cancer.

## Data availability

The data that support the findings of this study are available within the paper [and its supplementary information files] or from the corresponding author upon reasonable request. The proteomics data have been deposited to the ProteomeXchange Consortium via the PRIDE partner repository with the dataset identifier PXD016478 and will be made available upon publication.

## Conflict of interest

The authors declare no competing interests.

### Acknowledgements

The authors acknowledge Euro-BioImaging (www.eurobioimaging.eu) for providing access to imaging technologies and services via the Finnish Advanced Light Microscopy Node (Turku, Finland). We thank P. Laasola, J. Siivonen and Anja Gödicke for technical assistance, the Ivaska lab for critical reading of the manuscript and H. Hamidi for editing of the manuscript. We thank Virgilio Faila, Bernd Zobiak and Markku Saari for help with the microscopes and Dirk Mielenz, Aymelt Itzen and Peter Jackson for providing reagents. The Cell Imaging and Cytometry core (Turku Bioscience Centre, University of Turku and Åbo Akademi University and Biocenter Finland), the Euro-BioImaging Finnish Node, the Electron Microscopy Unit and Histocore at the University of Turku and the UMIF (UKE Microscopy Imaging Facility, Universitätsklinikum Hamburg, Eppendorf) are acknowledged for services, instrumentation and expertise. This study has been supported by DFG (VE 750/2-1) and Alexander von Humboldt-Foundation (S.V.), ProExzellenzia Hamburg and Academy of Finland (P.M.L.), an ERC CoG grant 615258 (J.I.); by grants and a fellowship from the National Health and Medical Research Council of Australia (grants APP1140064 and APP1150083 and fellowship APP1156489 to R.G.P.) and the Australian Research Council Centre of Excellence in Convergent Bio-Nano Science and Technology, CE140100036 (R.G.P.).

## AUTHOR CONTRIBUTIONS

Conceptualization, S.V. and J.I.; Methodology, P.M.L., N.Z.J., C.S., R.G., G.J., H.A., M.S., S.M., S.V., R.G.P., J.R., and J.I.; Formal Analysis, P.M.L., N.Z.J., C.S., R.G., G.J., P.K., R.E.K., L.O.F., S.M., M.S., S.V. R.G.P., J.R., and J.I.; Investigation, S.V., P.M.L. and J.I.; Resources, P.C., S.L., M.A., G.S. and A.D.; Writing – Original Draft, P.M.L. and J.I.; Writing – Review and Editing, S.V., P.M.L., G.J., N.Z.J., R.G.P. and J.I.; Supervision, P.M.L., S.V. and J.I.; Funding Acquisition, S.V., M.A., P.M.L., R.G.P. and J.I.

## Material and methods

### Cell culture, cell transfection and ECM coatings

MDA-MB-231 (triple negative human breast adenocarcinoma) cells were grown in Dulbecco’s Modified Eagle’s medium (DMEM) supplemented with 10 % fetal calf serum (FCS) and 1 % L-glutamine. Cells were purchased from the American Type Culture Collection (ATCC) and were routinely monitored for mycoplasma contamination. Mouse embryonic fibroblasts (MEFs) were cultured in DMEM supplemented with 20 % FCS, 1 % L-glutamine and 1 µg/ml puromycin. Human embryonic kidney 293 (HEK293) cells were cultured in DMEM supplemented with 20 % FCS and 1 % L-glutamine. Plasmids of interest were transfected using Lipofectamine 3000 (Thermo Fisher Scientific) or jetPRIME® (Polyplus transfection) according to the manufacturer’s instructions. The expression of proteins of interest was suppressed using 27 nM siRNA and lipofectamine siRNA Max (ThermoFisher Scientific) according to manufacturer’s instructions. The siRNA used as control (siCTRL) was Allstars negative control siRNA (Qiagen, Cat. No. 1027281). The siRNA oligonucleotides targeting swiprosin-1 were purchased from Sigma (siRNA#1 Cat. No. SASI_Hs01_00186848 and siRNA#2 Cat. No. SASI_Hs01_00186847). The siRNA oligonucleotides targeting Arf1 and IRSp53 were purchased from Qiagen (FlexiTube GeneSolution GS375 for Arf1; siArf1#1 Cat. No. SI02654470; siArf1#2 Cat. No. SI00299250; Qiagen FlexiTube IRSp53 siRNA#1 Cat. No. SI02637271; IRSp53 siRNA#2: Qiagen FlexiTube Cat. No. SI00087675). Coverslips were coated with either 10 µg/mL of fibronectin (FC010 Sigma), 12 µg/mL of laminin-1 (L4544 Sigma) or 300 µg/mL of collagen I from rat-tail (08-115 Sigma).

### Plasmids

Human Rab21 (aa 16-225) constructs were cloned into a Gateway destination vector pgLAP1 (Addgene plasmid #19702) to express Rab21 with an N-terminal GFP followed by a TEV cleavage site and an S-Tag in mammalian cells. As entry vector, we used pCR8/GW/Topo (Thermo Fisher, USA), in which human Rab21 was cloned by TOPO cloning. The Rab21 mutants Q76L (active) and T33N (inactive) were introduced into the entry clone by site-directed mutagenesis. Mouse full-length swiprosin-1, EF1-swiprosin-1 (deletion in the EF1 domain, aa 96-166) and EF2-swiprosin-1 (deletion at the EF2 domain, aa 134-159) were a kind gift from Dr. Dirk Mielenz (Division of Molecular Immunology, Nikolaus-Fiebiger-Centre, University of Erlangen-Nuremberg, Germany). The corresponding coding sequences were subcloned into the pEGFP-N1 backbone vector using the XhoI and EcoRI restriction sites. Full-length swiprosin-1 was also subcloned by TOPO cloning into the Gateway vector pCR8GWTopo. Then, a LR clonase II reaction was performed to shuttle full-length swiprosin-1 into pGEX-4T1. mScarlet-I swiprosin-1 was generated from the GFP-tagged construct by introducing the mScarlet-I sequence using the AgeI and MfeI restriction sites. Arf1-HA and Arf1-GFP were obtained from Addgene (plasmids #79409 and #49578). IRSp53-mcherry and IRSp53-GFP were generated by the Genome Biology Unit supported by HiLIFE and the Faculty of Medicine, University of Helsinki, and Biocenter Finland. pDEST-V1-Rab21 was generated by shuttling the human Rab21 sequence (pENTR201-hRab21,ORFeome Library; Genome Biology Unit supported by HiLIFE and the Faculty of Medicine, University of Helsinki, and Biocenter Finland) into the destination vector pDEST-V1-ORF (Addgene plasmid #73635). pDEST-swiprosin-1-V2 was generated by performing a clonase reaction between pCR8GWTopo-swiprosin-1 and pDEST-ORF-V2 (Addgene 73638). pEF.DEST51-mVenus was obtained from Addgene (plasmid #154899).

### Antibodies

The antibodies used in this work are listed in Supplementary Table 2.

### Generation of stable cell lines and SILAC-cell culture treatment

To generate stable GFP-Rab21 expressing mammalian cell lines, a FRT (Flp Recombination Target)-entry site was first introduced into MDA-MB-231 breast cancer cells using the Flp-In technology (Thermo Fisher, USA). Briefly, MDA-MB-231 cells were transfected with pFRT/lacZeo2 and stable clones were isolated using Zeocin as the selection marker (200 µg/ml). The resulting MDA-MB-231-FRT cell line was then co-transfected with the pgLAP1-Rab21 plasmid and pOG44 and stable clones were selected in 500 µg/ml Hygromycin-containing medium and tested for GFP-Rab21 expression. Stable MDA-MB-231-GFP-Rab21 cell lines were cultivated in heavy (Arg-10/Lys-8) or light SILAC-DMEM medium plus Hygromycin (200 µg/ml) for 12 days to ensure at least 10 replication cycles for efficient labelling. Hygromycin supplementation was omitted for the last two days before the co-IP experiment.

### Screening for Rab21 interaction partners by GFP pulldown

Each mass spectrometry experiment consisted of a mixture of a GFP-pulldown in active GFP-Rab21 expressing MDA-MB-231 cells (WT or active Rab21 Q76L mutant) grown in heavy medium and a GFP-pulldown in control cells (expressing GFP or inactive GFP-Rab21 T33N mutant) grown in light medium (forward experiment). In the reverse experiment, the heavy and light media were exchanged (label swap experiment). Briefly, co-IP samples were prepared as follows. Cells (two 15 cm dishes) were grown until 60-80% confluence, washed with ice cold PBSM (PBS + 5 mM MgCl_2_), harvested in PBSM, pooled and washed again. Cell pellets were resuspended in 600 µl lysis buffer LB (50 mM Tris, pH 7.5, 5 mM MgCl_2_, 150 mM KCl, 1.3 % n-beta-octyl-D-glucopyranoside, 10 % Glycerol, protease and phosphatase inhibitors and 500 µM GppNHp for GFP-active Rab21-expressing cells or 500 µM GDP for GFP-inactive Rab21-expressing cells) and lysed by douncing 40x in a tissue grinder (dounce homogenizer) and incubating for 20 min on ice. The insoluble fraction was removed by centrifugation at 18000x g and the supernatant was incubated with 20 µl GFP-trap agarose beads (Chromotek) for 60 min at 4°C by overhead rotation. Beads were then washed three times with 500 µl washing buffer WB (50 mM Tris, pH 7.5, 5 Mm MgCl_2_, 300 mM KCl, 10 % Glycerol), combining the heavy, GFP-active Rab21 IP with the light, GFP-inactive Rab21 IP (forward experiment) during the second washing step. Proteins were eluted from the beads in 100 µl of U/T buffer (6 M urea, + 2 M thiourea in 10 mM HEPES, pH 8.0) for 15 min with shaking at 1400 rpm at room temperature. The eluted proteins were collected and the process repeated a second time to maximize protein yield. Eluted proteins were precipitated by adding 70 µl 2.5 M Na-Acetate, pH 5.0, 1 µl GlycoBlue (Thermo Fisher, USA) and 1700 µl ethanol to the pooled elution fractions (200 µl) in a 2 ml tube. After an overnight incubation at 4°C, the precipitation mixture was centrifuged for 50 min at 20000x g and the resulting pellet dried for 15-20 min at 60-70°C. For in-solution protein digestion we followed standard procedures ^40^. Briefly, pellets were solubilized in U/T buffer, reduced with DTT, alkylated with iodoacetamide and digested by sequential addition of LysC (Wako) and Trypsin (Promega) overnight at room temperature. Peptides were desalted and stored on STAGE tips until LC-MS/MS analysis. Samples were analyzed using 240 min acetonitrile gradients on a 20 cm long 75 µm ID reversed phase column filled with 3 µm C18 beads (Dr. Maisch) using a Proxeon HPLC system (Thermo Fisher) coupled to the electrospray ion source of a Q Exactive Plus mass spectrometer (Thermo Fisher). The mass spectrometer was operated in the data dependent mode with a full scan (AGC target 3E6; R = 70,000) followed by up to ten MS2 scans (R = 17,500; maximal injection time 60 ms) and a dynamic exclusion for 30 sec. Raw files were analyzed using MaxQuant (version 1.5.2.8) using default parameters and searched against a Uniprot human protein database (2014-10). For visualization purposes, all the identified proteins from each experiment were plotted (Fig. 1A, Extended Data Fig. 1), where each spot corresponds to one identified protein by mass spectrometry. Each plot is representative of two independent experiments (forward and reverse, x- and y-axis), where every experiment consists of two independent IPs. The proteomics data have been deposited to the ProteomeXchange Consortium via the PRIDE ^41^ partner repository with the dataset identifier PXD016478. The complete dataset will be made available after publication. The Log10 transformed ratios of the proteins identified by mass spectrometry are also available from the Supplementary Information (Supplementary Table 1).

### Immunoprecipitations, BICAP and immunoblotting

MDA-MB-231 or HEK293 cells expressing GFP-tagged or split Venus-tagged (V1 or V2) proteins (one 10 cm dish per condition) were washed with cold PBS, harvested in PBS and pelleted. The cell pellet was resuspended in 200 µl of IP-lysis buffer (40 mM Hepes-NaOH, 75 mM NaCl, 2 mM EDTA, 1% NP40, protease and phosphatase inhibitors) and incubated at +4 °C for 30 min, followed by centrifugation (10,000x g for 10 min, +4 °C). 20 µl of the supernatant was kept aside as the lysate control. The remainder of the supernatant was incubated with GFP-Trap beads (ChromoTek; gtak-20), which bind to both GFP and to complemented Venus (V1+V2), for 1 h at 4 °C. Finally, immunoprecipitated complexes were washed three times with wash-buffer (20 mM Tris-HCl pH 7.5, 150 mM NaCl, 1 % NP-40) and denatured for 5 min at 95°C in reducing Laemmli buffer before SDS-PAGE analysis under denaturing conditions (4–20 % Mini-PROTEAN TGX Gels). The proteins were then transferred to nitrocellulose membranes (Bio-Rad Laboratories) before blocking with blocking buffer (Thermo, StartingBlock (PBS) blocking, #37538) and PBS (1:1 ratio). The membranes were incubated with primary antibodies diluted in blocking buffer overnight at 4°C. Following this step, membranes were washed three times with TBST and incubated with fluorophore-conjugated secondary antibodies (LI-COR) diluted (1:10,000) in blocking buffer at room temperature for 1 hour. Membranes were scanned using an infrared imaging system (Odyssey; LI-COR Biosciences).

### Protein purification

For producing recombinant GST-tagged Rab21 16-225 and swiprosin-1, pGEX-4T1-Rab21 16-225 or pGEX-4T1-swiprosin-1, BL21 Rosetta transformed cells were grown and induced with 250 µM IPTG at OD 0.5-0.8 at 22°C in LB media overnight. Cells were then lysed and resuspended in 20 mM Tris, pH 7,5; 5 mM MgCl_2_, 300 mM NaCl, 3 mM DTT. The desired GST-tagged protein was purified from this suspension with a gravity GST-column. In the case of GST-Rab21, elution of the protein was followed by thrombin-cleavage and gel filtration in 20 mM Tris, pH 7.5, 5 mM MgCl_2_, 50 mM NaCl, 3 mM DTT buffer using a Superdex S75 16/60 column attached to a GST-column to bind cleaved GST. Fractions containing the monomeric proteins were pooled, concentrated via ultrafiltration (Amicon Ultra, 15 mL, 10,000 MWCO) and flash-frozen in liquid nitrogen for long-term storage at -80°C.

### Nucleotide loading and GST pulldowns

To load Rab21 with either GppNHp or GDP, 200 µM of recombinant Rab21 or GST-Rab21 were incubated with 10 mM EDTA and a 25x excess of nucleotide (5 mM GppNHp or GDP) in exchange buffer (20 mM Tris pH 7.5, 2.5 mM MgCl_2_, 50 mM NaCl and 3 mM DTT) for 1 h at 25°C. The EDTA-based exchange reaction was then stopped by the addition of 40 mM MgCl_2_ and incubation for 15 min on ice. The buffer was then exchanged with measuring buffer (20 mM Tris, pH 7.5, 5 mM MgCl_2_, 50 mM NaCl and 3 mM DTT) to reach the desired protein concentration using 10 kDa ultrafiltration devices (Amicon). Equal amounts of nucleotide compared to the protein amount was added to ensure complete loading of Rab21 with the desired nucleotide. GST pulldowns using purified GST fusion proteins and recombinant Rab21 bound to a non-hydrolysable form of GTP (GppNHp), GDP or no nucleotide (in the presence of EDTA) were performed as follows. 50 µg of GST-swiprosin-1 or GST-Appl1 was incubated with 200 µg of recombinant Rab21 for 30 min before incubation with Glutathione Sepharose beads for an additional 30 min. The proteins bound to the beads were divided in 2 fractions and separated by SDS-PAGE electrophoresis, followed by either Coomassie staining or immunoblotting with anti-Rab21 antibody. Pulldowns were performed three times.

### SIM microscopy and co-localization analysis

Cells growing on collagen-1-coated dishes were fixed with 2 % formaldehyde, permeabilized with 0.3 % Triton X-100, blocked with 10 % horse serum and incubated with antibodies against the indicated endogenous proteins. This was followed by incubation with fluorophore-labelled secondary antibodies (Alexa-568 or Alexa-488 anti-mouse or anti-rabbit, Life Technologies). For visualization of swiprosin-1-containing invaginations, cells expressing mScarlet-I-swiprosin-1 were incubated at 4°C for 30 min before fixation. Cells were imaged with an OMX DeltaVision and spots co-localization analysis was performed in Z stacks of the cells using the plugin ComDet in ImageJ (https://imagej.net/Spots_colocalization_(ComDet)), which allowed for a better detection of the invaginations since it ignores inhomogeneous cytoplasmic background. Using this plugin, we pinpointed the mScarlet-I-swiprosin-1 spots (pixel size above 5), and analyzed co-localization, based on proximity (pixel distance 4), with spots from the second channel stained for the protein of interest. Then, the ratio of colocalization with mScarlet-I-swiprosin-1 was calculated as a percentage of co-localized spots per cell. At least 30 cells per condition were imaged and analyzed. Co-localization plots show mean ± 95 % CI. Statistical significance was calculated compared to the control condition. Representative rendered images of the invaginations (x-z plane) were visualized with IMARIS software (Oxford Instruments).

### Proximity Ligation Assay (PLA)

MDA-MB-231 cells growing on coverslips were fixed, washed twice with PBS and permeabilized with 0.3 % Triton-X-100 in PBS for 15 min at room temperature. Cells were stained using anti-swiprosin-1 (1:100) and anti-Rab21 (1:50) primary antibodies diluted in 5 % horse serum for 1 h at room temperature. Proximity ligation was performed according to the manufacturer’s instructions (Duolink in situ PLA, Sigma-Aldrich). Interaction between swiprosin-1 and Rab21 in cells was detected using confocal microscopy (Leica TCS SP5, 63×1.4 Apo oil objective) and the number of PLA spots per 1000 µm^3^ was quantified using IMARIS software.

### BiFC and TIRF microscopy

MDA-MB-231 cells growing in glass-bottom dishes were co-transfected with split Venus constructs (pDEST-V1-Rab21 and pDEST-swiprosin-1-V2) and imaged 30 h after transfection. Imaging was performed using a DeltaVision OMX v4 (GE Healthcare Life Sciences) fitted with an Olympus APO N 60x Oil TIRF objective lens, 1.49 NA used in TIRF illumination mode. Emitted light was collected on a front illuminated pco.edge sCMOS (pixel size 6.5 mm, readout speed 95 MHz; PCO AG) controlled by SoftWorx. The TIRF angle for all channels was maintained at 83.5°.Images were taken every 500 ms for 2 min, at 37 °C in presence of 5 % CO_2_.

### DNA paint

For two-color single molecule localization microscopy (SMLM) (Extended Data Fig. 4), we used DNA-PAINT 23. Cells over-expressing EFHD2-GFP and ARF1-HA were fixed and labeled using primary antibodies against GFP (Abcam ab1218) and HA-tag (Cell Signaling 3724), respectively. Cells were then stained with appropriate secondary antibodies coupled to PAINT DNA handles (Ultivue). Imaging was performed using a DeltaVision OMX v4 (GE Healthcare Life Sciences) fitted with an Olympus APO N 60x Oil TIRF objective lens, 1.49 NA used in TIRF illumination mode. Emitted light was collected on a front illuminated pco.edge sCMOS (pixel size 6.5 mm, readout speed 95 MHz; PCO AG) controlled by SoftWorx. First, a TIRF image of EFHD2-GFP was acquired, followed by the DNA-paint acquisitions. DNA paint imaging was done sequentially first for EFHD2-GFP (10 000 frames, 50 ms) and for ARF1 (10 000 frames, 100 ms) in PAINT buffer (10 mM Tris-HCl, 100 mM NaCl, 0.05 % Tween-20, pH 7.5) supplemented with 0.5 nM of the corresponding PAINT imager strands coupled to AlexaFluor 647 (EFHD2-GFP) or to AlexaFluor 568 (ARF1). For both conditions, full laser power was used and the beam concentrator enabled. No cross talk between the channels was observed. The ThunderSTORM ^42^ ImageJ plugin ^43^, with the Phasor-based localization 2D method ^44^, was used for the localization of single fluorophores. After filtering out localizations to reject too low photon counts, translational shifts were corrected by autocorrelation. Image reconstructions were performed using the ThunderSTORM ImageJ plugin.

### Uptake/endocytosis assays

MDA-MB-231 cells were grown on collagen-1 coated-dishes or coverslips unless stated otherwise. For integrin endocytosis assays, surface integrins were labelled with an antibody that recognizes the active conformation of β1-integrins (12G10) at 4 °C followed by incubation at 37 °C for 15 min unless stated otherwise. The antibody still left on the surface was washed away with acid (0.2 M Acetic acid, 0.5 M NaCl pH 2.5). The cells were subsequently fixed with 2 % formaldehyde, permeabilized with 0.05 % saponin and incubated with a fluorescent secondary antibody to visualize and quantify the amount of internalized integrins. Several fields were randomly imaged with identical microscope settings using a 3i (Intelligent Imaging Innovations, 3i Inc) Marianas Spinning disk confocal microscope with a Yokogawa CSU-W1 scanner and a back illuminated 10 MHz EMCDD camera (Photometrics Evolve) using 63x/1.4 oil objective. Quantification of endocytosed integrins was performed on 3D-projections of the cells using IMARIS software with the spots detection function. The intensity sum of all the vesicles per cell was divided by the volume of that cell. All the intensity values were then normalized to the average of all the cells in the control condition (siCTRL). A similar procedure was followed for the uptake of MHCI (SAB4700637 Sigma) and 9EG7 (active β1-integrin). For AlexaFluor 568-labelled transferrin (Transferrin-AF568, Thermo Fisher T23365) and 10 kDa dextran uptake experiments, cells were treated with 1 mg/mL of transferrin, 10 kDa dextran-TMR (D1816, Invitrogen) or 10 kDa amino dextran (Molecular Probes) conjugated to FITC during the 15 min incubation at 37°C. In the case of the double uptake experiments, cells were previously labelled with 12G10-AF488 antibody (Abcam, ab202641) at 4°C as described above. After the internalization step at at 37°C, the remaining fluorescently labelled molecules at the cell surface were removed with an acid wash, followed by fixation, labelling of plasma membrane with WGA lectin and imaging as described above.

### Electron microscopy and Apex labelling

Apex labeling of GFP-swiprosin-1 for electron microscopy was performed as previously described 28. Briefly, MEFs were transfected with GFP-swiprosin-1 and GBP-Apex (Addgene plasmid 67651) constructs using a Neon transfection system (as per manufacturer’s instructions) and plated into 35 mm tissue culture dishes. 24 h post transfection, cells were incubated on ice for 30 min, then fixed in 2.5 % glutaraldehyde, washed 3x in cacodylate buffer and once in 1 mg/ml 3,3-diaminobenzidine (DAB; Sigma-Aldrich) solution in cacodylate for 5 min. Cells were then subjected to the DAB reaction in the presence of H2O2 for 30 min and, stained with 1 % osmium tetroxide for 2 min. The cells were processed in situ with serial dehydration in increasing percentages of ethanol, then serial infiltration with LX112 resin in a Pelco Biowave microwave before polymerizing overnight at 60°C. Ultra-thin (60 nm) sections were cut on a Leica UC6 microtome parallel to the culture dish and micrographs acquired using a Jeol 1011 transmission electron microscope.

### Vesicle tracking

For Rab21 vesicle tracking, MDA-MB-231 cells stably expressing GFP-Rab21 were transfected with one of two different siRNA sequences to deplete the levels of swiprosin-1. The movement of the Rab21 vesicles was followed for 2 min using TIRF microscopy (Visitron SD-TIRF Nikon Eclipse TiE with a 60x Olympus TIRF oil objective, NA: 1.49). The average speed of the vesicles was measured in each cell. At least 10 cells per condition were analyzed and data from 3 independent experiments was quantified.

### Focal adhesion analysis

Cells were plated on dishes coated with the indicated extracellular matrix component (collagen I, laminin-1 or fibronectin), fixed and immunostained with an antibody that recognizes the focal adhesion component vinculin, present in mature adhesions. At least 6 cells per condition per experiment were imaged and analyzed (18 cells per condition in total from 3 independent experiments). Vinculin-positive focal adhesions were detected from an image mask created using the ImageJ software following background subtraction and setting of median Gaussian filter (3.0). The number of focal adhesions (detected particles) and the total cell area occupied by focal adhesions per cell was quantified from the image mask. Plots show mean ± 95 % CI.

### Adhesion dynamics

For adhesion dynamics studies, MDA-MB-231 cells transiently expressing GFP-paxillin were transfected with one of two different siRNA sequences to deplete the levels of swiprosin-1. GFP-paxillin was imaged for 120 min at the TIRF plane with a Deltavision OMX using a 63x objective. Cells were imaged every 1 min, at 37°C in presence of 5 % CO2, using multiposition capabilities. Focal adhesion dynamics were then analyzed using the Focal Adhesion Analysis Server ^45^. Only focal adhesions with a lifetime minimum of 10 frames were analyzed, and the focal adhesions that were assembled and disassembled during the course of imaging were used for measuring focal adhesion kinetics.

### Migration assay

After siRNA transfection, MDA-MB-231 cells were seeded into ibidi 2-well culture inserts placed in ibidi µ-dishes and allowed to become confluent. Prior to imaging, the culture inserts were carefully removed with forceps and the cells were washed twice with PBS. Next, warm medium was added to the cells after which live imaging was started immediately (37 °C, 5 % CO_2_). Cells were imaged using a Nikon Eclipse Ti2-E microscope with a 10x objective for 24 h with a 20 minute imaging interval. Data was quantified using Fiji (ImageJ) by measuring the area of the closing gap between the cells at 0, 6, 8, 12, 16, and 18 h.

### Invasion assay

200 µl of PureCol EZ gel (#5074; Advanced Biomatrix) was allowed to polymerize in 8 µm inserts (662638; Greiner Bio-One) for 1 h at 37 °C. Inserts were then inverted and 100 µl of cell suspension (50000 cells) were seeded onto the outer face of the insert. Cells were allowed to adhere at 37 °C for 3 h. Inserts were then dipped sequentially into PBS and placed in serum-free medium. Medium supplemented with 10 % of FCS and 20 ng/ml of EGF was placed on top of the matrix and cells were allowed to invade the matrix for 72 h. Cells were then fixed using 4 % PFA for 2 h, permeabilized in 0.3 % Triton-X 100 for 1 h at room temperature, and stained overnight at 4 °C using Alexa Fluor 488 phalloidin. Invading cells were imaged using a confocal microscope (LSM880; Zeiss). Invasion was quantified using the area calculator plugin in ImageJ, measuring the fluorescence intensity of cells invading 60 µm or more and expressing this as a percentage of the fluorescence intensity of all cells within the matrix.

### Statistical analysis for in vitro samples

The GraphPad Prism software and two-tailed Student’s t test (paired or unpaired, as appropriate) was used for statistical analysis. When data were not normally distributed, a MannWhitney test was used. P values<0.05 are shown in graphs. For all graphs, ns = not significant.

### Breast cancer tissue microarrays

The study was approved by the Hospital District of Southwest Finland and Turku University Hospital (decision T012/015/19) and the use of tissue samples was approved by the Scientific Steering Group of Auria Biobank (decision AB19-4522). The study population consisted of 243 patients with breast cancer diagnosed and treated in Turku University Hospital in 1998-2013. All patients were treated with surgical resection or mastectomy and the archived formalin fixed, paraffin-embedded tumor samples were used to form tissue microarrays (TMAs) that were prepared similarly as previously described ^46^. Briefly, the TMAs were prepared by punching a representative site of paraffin block of each tumor with either a 1 mm or a 1.5 mm diameter cylinder and using an automated tissue arrayer (TMA Grand Master, 3DHISTECH Ltd., Budapest, Hungary). The cohort consisted of 149 patients with triple negative and 89 patients with HER2-positive breast cancer diagnosed using the WHO classification criteria of tumors of the breast at the time of sampling. In addition, 4 patients with hormone receptor positive and 1 with non-neoplastic breast cancer were included in the TMAs. The cores were available from 225 tumor centers, 121 tumor borders, 26 lymph node metastases and 127 tumor areas with inflammatory infiltrate. All relevant medical records of the patients were reviewed and information on tumor size, histological grade, hormone receptor status, Her2-oncogene, proliferation marker Ki-67 and axillary lymph node status were gathered. The follow-up time was until 31st March 2020 and the range of follow-up varied from 1 month to 22 years 3 months (mean 10 years 2 months).

### Immunohistochemistry

Immunohistochemistry was performed on TMAs comprising one or two tissue cores from each tumor site of each patient. The tissue samples were cut into 4µm sections, deparaffinized and rehydrated with standard procedures. Heat-mediated antigen retrieval was done for all samples in citrate buffer (pH 6) in a pressure cooker (Decloaking chamber, Biocare Medical NxGen) for 20 min. Sections were stained in a semi-automatic Labvision autostainer (Thermo-Fisher Scientific), where they were washed with washing buffer (0.05M Tris-HCl pH 7.6, 0.05 % Tween 20) and the endogenous enzymes were blocked with 3 % H2O2 Tris-HCl for 10 min. This was followed by a blocking step using Normal Antibody Diluent (NABD; Immunologic, BD09-125), incubation with the primary antibody (anti-swiprosin-1, Atlas Antibodies, diluted 1:200) for 1h, followed by washes and incubation with the secondary antibody (Goat anti-rabbit HRP, Immunologic DPVB110HRP) for 30min. Samples were then washed and incubated with the DAB solution (Bright DAB, Immunologic BS04-110) for 10 min. After counterstain with Mayer’s HTX, slides were dehydrated, cleared in xylene and mounted with Pertex. Antibody specificity was validated on agarose-embedded cell pellets post siRNA transfection (siCTRL, Swip1 siRNA 1 and Swip1 siRNA#2; Extended Data Fig. 8A). For each tumor sample, the percentage of cells with immunopositive signal (0-100%) in cytoplasm and in plasma membrane were scored. Samples with more than 80 % of cells exhibiting positive staining were considered as having high swiprosin-1, those with less than 30 % of cells as low swiprosin-1 and samples in between were considered to have medium swiprosin-1 levels.

### Statistical analysis for clinical samples

The statistical analysis was performed with SPSS Statistics 26 (IBM Corp., NY, USA). Two-tailed p values below 0.05 were considered statistically significant. The clinical parameters (age, tumor size and ki67) across swiprosin-1 categories (<100 and 100, or <80 and 80) were evaluated with independent-samples Mann-Whitney U tests. The categorical parameters (grade and lymph node metastasis status) were compared with Chi Squared tests or Fisher’s exact tests across swiprosin-1 categories. Related-samples Wilcoxon signed rank test was used for paired comparisons of tumor centers and other core types. Patients with missing data were censored from the paired comparisons. Overall survival was compared between low and high percentage of immunopositive cells in the tumor center samples of Her2+ and TNBC patients using Kaplan-Meier plots and log-rank tests: the samples were divided into the following groups: <100 % or 100 % in the case of the cytoplasmic signal and <80 % or 80 % in the case of membranal swiprosin-1 signal. The Cox proportional hazards regression analysis was used to assess the hazard ratio of tumor centers with high vs. lower swiprosin-1 immunopositivity. In the case of non-proportional hazards, the weighted estimation of Cox regression was used ^47^. The adjusted Cox analysis was performed relative to the clinical prognostic features. The analysis was performed with R Software for Statistical Computing (version 3.6.2, www.r-project.org) and survival (version 3.1.8) and coxphw (version 4.0.2) packages. Survival differences were quantified as hazard ratios (HRs) with 95 % confidence intervals (CI).

## Bibliography

1. Kaksonen, M. Roux, A. Mechanisms of clathrin-mediated endocytosis. Nat. Rev. Mol. Cell Biol.s 19, 313–326 (2018).

2. Mettlen, M., Chen, P.-H., Srinivasan, S., Danuser, G. Schmid, S. L. Regulation of Clathrin-Mediated Endocytosis. Annu. Rev. Biochem. 87, 871–896 (2018).

3. Thottacherry, J. J., Sathe, M., Prabhakara, C. Mayor, S. Spoiled for Choice: Diverse Endocytic Pathways Function at the Cell Surface. Annu. Rev. Cell Dev. Biol. 35, 55–84 (2019).

4. Sanger, A., Hirst, J., Davies, A. K. Robinson, M. S. Adaptor protein complexes and disease at a glance. J. Cell Sci. 132, (2019).

5. Mayor, S., Parton, R. G. Donaldson, J. G. Clathrin-independent pathways of endocytosis. Cold Spring Harb. Perspect. Biol. 6, (2014).

6. Maldonado-Báez, L., Williamson, C. Donaldson, J. G. Clathrin-independent endocytosis: a cargo-centric view. Exp. Cell Res. 319, 2759–2769 (2013).

7. Ferreira, A. P. A. Boucrot, E. Mechanisms of Carrier Formation during Clathrin-Independent Endocytosis. Trends Cell Biol. 28, 188–200 (2018).

8. Kirkham, M. et al. Ultrastructural identification of uncoated caveolin-independent early endocytic vehicles. J. Cell Biol. 168, 465–476 (2005).

9. Howes, M. T. et al. Clathrin-independent carriers form a high capacity endocytic sorting system at the leading edge of migrating cells. J. Cell Biol. 190, 675–691 (2010).

10. Sathe, M. et al. Small GTPases and BAR domain proteins regulate branched actin polymerisation for clathrin and dynamin-independent endocytosis. Nat. Commun. 9, 1835 (2018).

11. Pellinen, T. et al. Small GTPase Rab21 regulates cell adhesion and controls endosomal traffic of beta1-integrins. J. Cell Biol. 173, 767–780 (2006).

12. Pellinen, T. et al. Integrin trafficking regulated by Rab21 is necessary for cytokinesis. Dev. Cell 15, 371–385 (2008).

13. Högnäs, G. et al. Cytokinesis failure due to derailed integrin traffic induces aneuploidy and oncogenic transformation in vitro and in vivo. Oncogene 31, 3597–3606 (2012).

14. Alanko, J. et al. Integrin endosomal signalling suppresses anoikis. Nat. Cell Biol. 17, 1412–1421 (2015).

15. Moreno-Layseca, P., Icha, J., Hamidi, H. Ivaska, J. Integrin trafficking in cells and tissues. Nat. Cell Biol. 21, 122–132 (2019).

16. Meyer, K. Selbach, M. Quantitative affinity purification mass spectrometry: a versatile technology to study protein-protein interactions. Front. Genet. 6, 237 (2015).

17. Hubner, N. C. et al. Quantitative proteomics combined with BAC TransgeneOmics reveals in vivo protein interactions. J. Cell Biol. 189, 739–754 (2010).

18. Jean, S. Kiger, A. A. RAB21 Activity Assay Using GST-fused APPL1. Bio-Protoc. 6, (2016).

19. Zhu, G. et al. Structure of the APPL1 BAR-PH domain and characterization of its interaction with Rab5. EMBO J. 26, 3484–3493 (2007).

20. Croucher, D. R. et al. Bimolecular complementation affinity purification (BiCAP) reveals dimer-specific protein interactions for ERBB2 dimers. Sci. Signal. 9, ra69 (2016).

21. Kumari, S. Mayor, S. ARF1 is directly involved in dynamin-independent endocytosis. Nat. Cell Biol. 10, 30–41 (2008).

22. Arjonen, A., Alanko, J., Veltel, S. Ivaska, J. Distinct recycling of active and inactive β1 integrins. Traffic Cph. Den. 13, 610–625 (2012).

23. Schnitzbauer, J., Strauss, M. T., Schlichthaerle, T., Schueder, F. Jungmann, R. Super-resolution microscopy with DNA-PAINT. Nat. Protoc. 12, 1198–1228 (2017).

24. Caldieri, G. et al. Reticulon 3-dependent ER-PM contact sites control EGFR nonclathrin endocytosis. Science 356, 617–624 (2017).

25. McMahon, H. T. Boucrot, E. Molecular mechanism and physiological functions of clathrin-mediated endocytosis. Nat. Rev. Mol. Cell Biol. 12, 517–533 (2011).

26. Hemalatha, A. Mayor, S. Recent advances in clathrin-independent endocytosis. F1000Research 8, (2019).

27. Park, K. R. et al. Structural implications of Ca2+-dependent actin-bundling function of human EFhd2/Swiprosin-1. Sci. Rep. 6, (2016).

28. Ariotti, N. et al. Modular Detection of GFP-Labeled Proteins for Rapid Screening by Electron Mi-croscopy in Cells and Organisms. Dev. Cell 35, 513–525 (2015).

29. Paul, N. R., Jacquemet, G. Caswell, P. T. Endocytic Trafficking of Integrins in Cell Migration. Curr. Biol. CB 25, R1092–1105 (2015).

30. Disanza, A. et al. CDC42 switches IRSp53 from inhibition of actin growth to elongation by clustering of VASP. EMBO J. 32, 2735–2750 (2013).

31. Krugmann, S. et al. Cdc42 induces filopodia by promoting the formation of an IRSp53:Mena complex. Curr. Biol. CB 11, 1645–1655 (2001).

32. Lim, K. B. et al. The Cdc42 effector IRSp53 generates filopodia by coupling membrane protrusion with actin dynamics. J. Biol. Chem. 283, 20454–20472 (2008).

33. Pipathsouk, A. et al. WAVE complex self-organization templates lamellipodial formation. bioRxiv 836585 (2019) doi:10.1101/836585.

34. Koronakis, V. et al. WAVE regulatory complex activation by cooperating GTPases Arf and Rac1. Proc. Natl. Acad. Sci. U. S. A. 108, 14449–14454 (2011).

35. Schlienger, S., Ramirez, R. A. M. Claing, A. ARF1 regulates adhesion of MDA-MB-231 invasive breast cancer cells through formation of focal adhesions. Cell. Signal. 27, 403–415 (2015).

36. Norman, J. C. et al. ARF1 mediates paxillin recruitment to focal adhesions and potentiates Rho-stimulated stress fiber formation in intact and permeabilized Swiss 3T3 fibroblasts. J. Cell Biol. 143, 1981–1995 (1998).

37. Lundmark, R. et al. The GTPase-activating protein GRAF1 regulates the CLIC/GEEC endocytic pathway. Curr. Biol. CB 18, 1802–1808 (2008).

38. Hornbruch-Freitag, C., Griemert, B., Buttgereit, D. Renkawitz-Pohl, R. Drosophila Swiprosin-1/EFHD2 accumulates at the prefusion complex stage during Drosophila myoblast fusion. J. Cell Sci. 124, 3266–3278 (2011).

39. Del Olmo, T. et al. APEX2-mediated RAB proximity labeling identifies a role for RAB21 in clathrin-independent cargo sorting. EMBO Rep. 20, (2019).

40. Paul, F. E., Hosp, F. Selbach, M. Analyzing protein-protein interactions by quantitative mass spectrometry. Methods San Diego Calif 54, 387–395 (2011).

41. Perez-Riverol, Y. et al. The PRIDE database and related tools and resources in 2019: improving support for quantification data. Nucleic Acids Res. 47, D442–D450 (2019).

42. Ovesný, M., Křrížek, P., Borkovec, J., Svindrych, Z. Hagen, G. M. ThunderSTORM: a comprehensive ImageJ plug-in for PALM and STORM data analysis and super-resolution imaging. Bioinforma. Oxf. Engl. 30, 2389–2390 (2014).

43. Schindelin, J. et al. Fiji: an open-source platform for biological-image analysis. Nat. Methods 9, 676–682 (2012).

44. Martens, K. J. A., Bader, A. N., Baas, S., Rieger, B. Hohlbein, J. Phasor based single-molecule localization microscopy in 3D (pSMLM-3D): An algorithm for MHz localization rates using standard CPUs. J. Chem. Phys. 148, 123311 (2018).

45. Berginski, M. E. Gomez, S. M. The Focal Adhesion Analysis Server: a web tool for analyzing focal adhesion dynamics. F1000Research 2, 68 (2013).

46. Heuser, V. D. et al. Formin Proteins FHOD1 and INF2 in Triple-Negative Breast Cancer: Association With Basal Markers and Functional Activities. Breast Cancer Basic Clin. Res. 12, 1178223418792247 (2018).

47. Dunkler, D., Ploner, M., Schemper, M. Heinze, G. Weighted Cox Regression Using the R Package coxphw. J. Stat. Softw. 84, 1–26 (2018).

