## Extended data figures and supplementary information for "Cargo-specific recruitment in clathrin and dynamin-independent endocytosis"

Extended Data Figures 1-8 and associated legends

Supplementary tables 1-2, supplementary videos and video legends 1-5,

### Extended data figures and figure legends

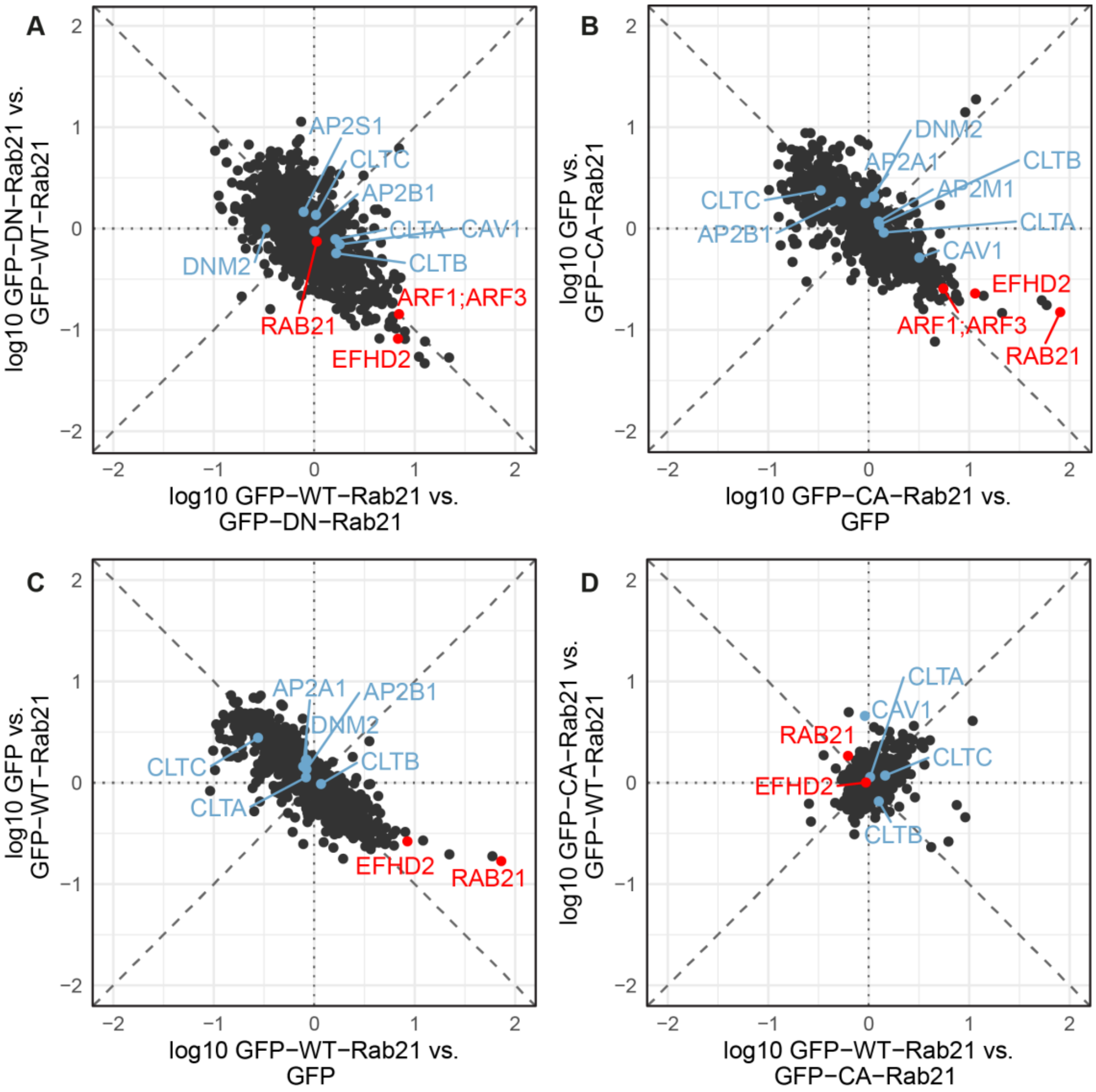


**Extended Data Fig. 1. Swiprosin-1 (EFHD2, Swip1) is an interactor for Rab21.**
SILAC-MS analysis of Rab21 interactors from MDA-MB-231 cells expressing GFP-WT-Rab21 vs. GFP-DN-Rab21 (T31N inactive GDP-bound/nucleotide-free mutant) (A); GFP-CA-Rab21 (Q76L constitutively active GTP-bound mutant) vs. GFP (B); GFP-WT-Rab21 vs. GFP (C) or GFP-WT-Rab21 vs. GFP-CA-Rab21 (D). Each spot corresponds to one identified protein by mass spectrometry. Plots are representative of two independent experiments (forward and reverse, x- and y-axis), where every experiment consists of two independent affinity purifications. For example, in (A), swip1 is enriched by approx. 12.2x and 6.8x, and Arf1 is enriched 7.0x and 7.0x in the GFP-WT-Rab21 fraction compared to GFP-DN-Rab21 (forward and reverse experiment, respectively). In (B), swip1 is enriched about 4.4x and 11.5x, and Arf1 is enriched 3.9x and 5.5x. Swip1 was strongly enriched in the GFP-WT-Rab21 vs. GFP-DN-Rab21 (A) and GFP-CA-Rab21 and GFP-WT-Rab21 fractions compared to GFP fractions (B-C), and it was equally enriched in the GFP-CA-Rab21 compared to the GFP-WT-Rab21 fraction (D). Clathrin (CLTA, CLTB, CLTC), AP2 (AP2A1, AP2B1, AP2M1, AP2S1), caveolin (CAV1) and dynamin II (DNM2) are not strongly enriched in any fraction.

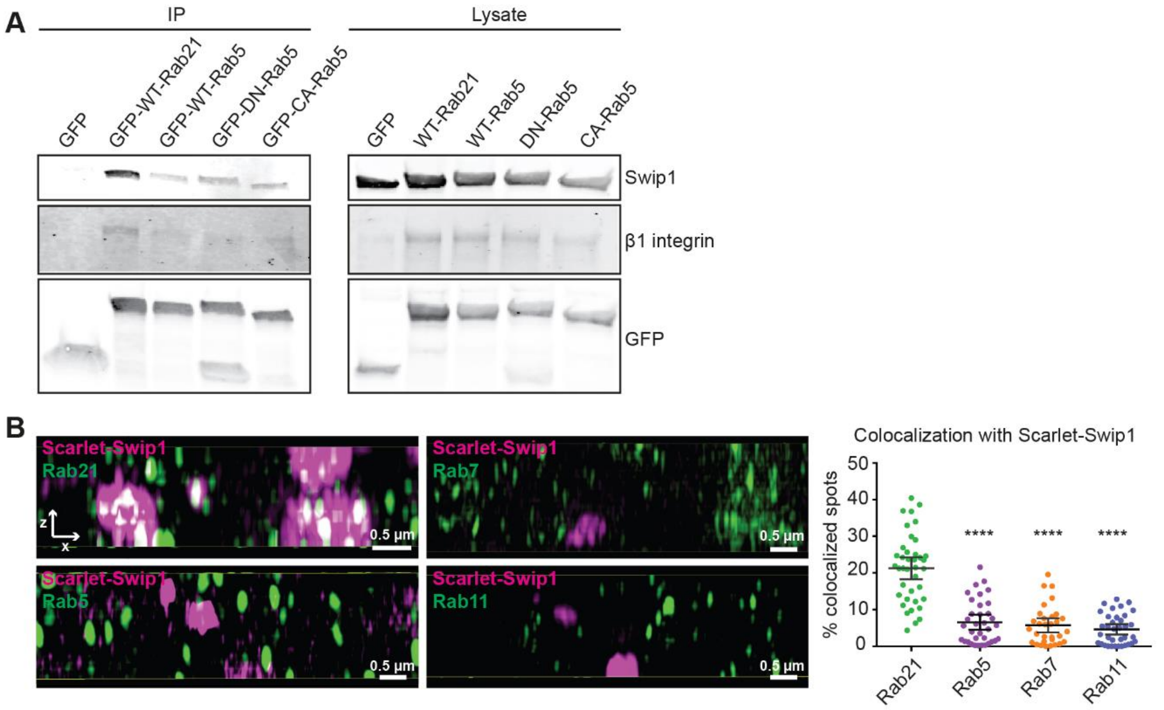


**Extended Data Fig. 2. Swip1 binds preferentially to Rab21 compared to other Rabs.**
(A) Representative immunoblots of GFP-TRAP pulldowns from MDA-MB-231 cells transfected with GFP, GFP-WT-Rab21, GFP-WT-Rab5, GFP-DN-Rab5 or GFP-CA-Rab5 stained for endogenous swip1 and β1 integrin. n = 3 independent experiments.
(B) SIM x-z projections of MDA-MB-231 cells expressing mScarlet-I-Swip1 and immunostained for endogenous Rab proteins and quantification of Rab protein co-localization with mScarlet-I-Swip1. Each dot represents the co-localization ratio in one cell. n = 39 cells for Rab21, 35 for Rab5, 31 for Rab7, 36 for Rab11. Data are mean ± 95 % CI, **** *P* < 0.0001, Mann Whitney test, n = 3.

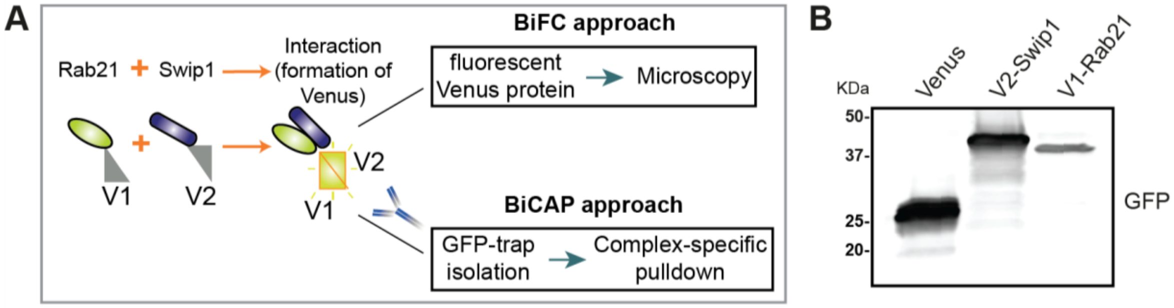


**Extended Data Fig. 3. BiFC/BiCAP approaches to assess the interaction between Rab21 and swip1.** (A) Cartoon of the BiFC/BiCAP approach used to image (BiFC) and biochemically detect (BiCAP) the interactions between Rab21 and swip1 in live cells and from cell lysate. (B) MDA-MB-231 cells were transfected with the indicated constructs and lysed. Proteins expressed in the transfected cells were detected by immunoblotting. Immunoblot shows the size of the bands detected by a polyclonal anti-GFP antibody.

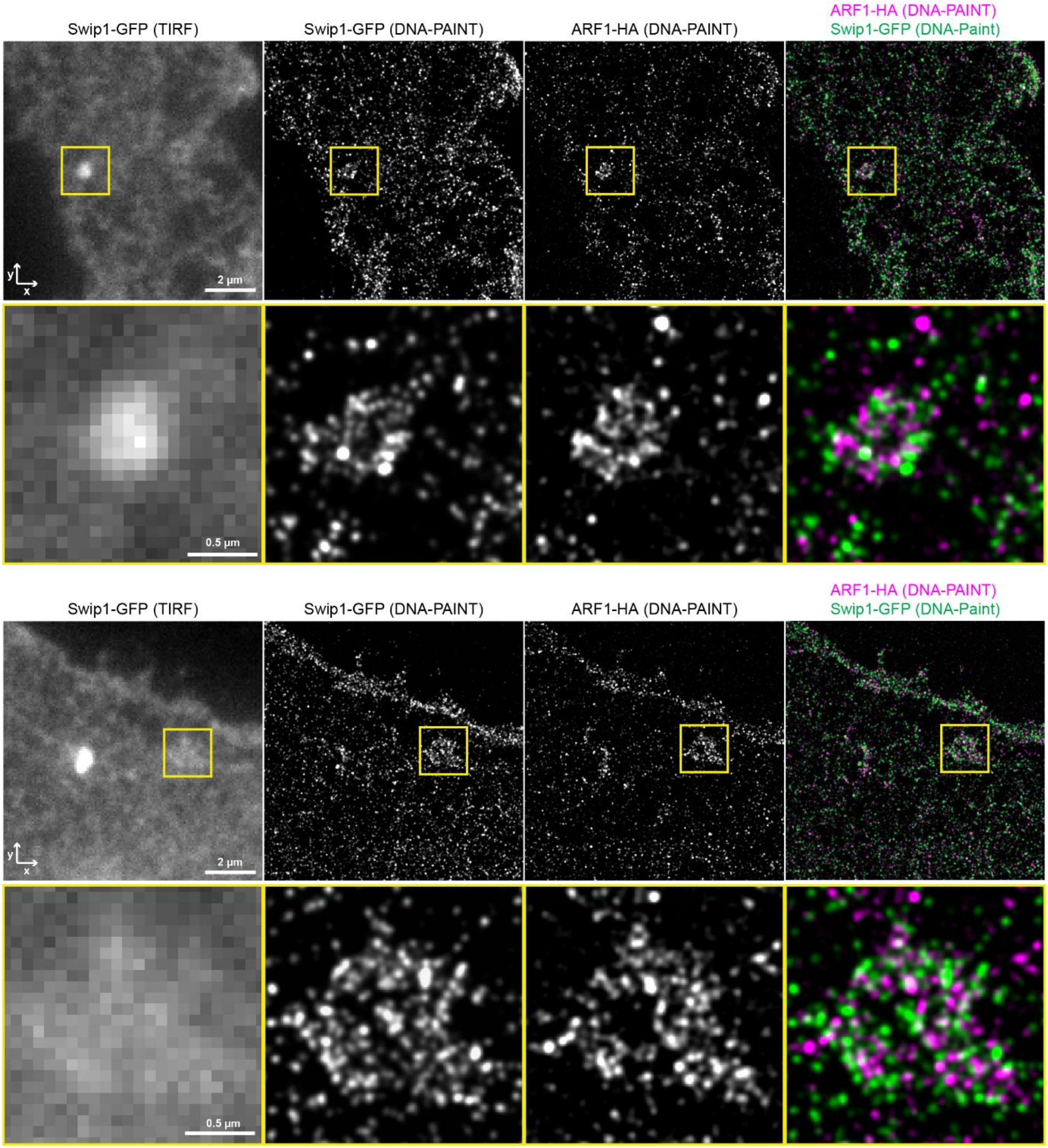


**Extended Data Fig. 4. GFP-swip1 co-localizes with Arf1-HA at the TIRF plane.**
MDA-MB-231 cells were co-transfected with GFP-swip1 and Arf1-HA constructs, fixed, stained with probes for DNA paint and imaged using SMLM. Examples of the structures formed by GFP-swip1 at the proximity of the ECM interphase are shown.

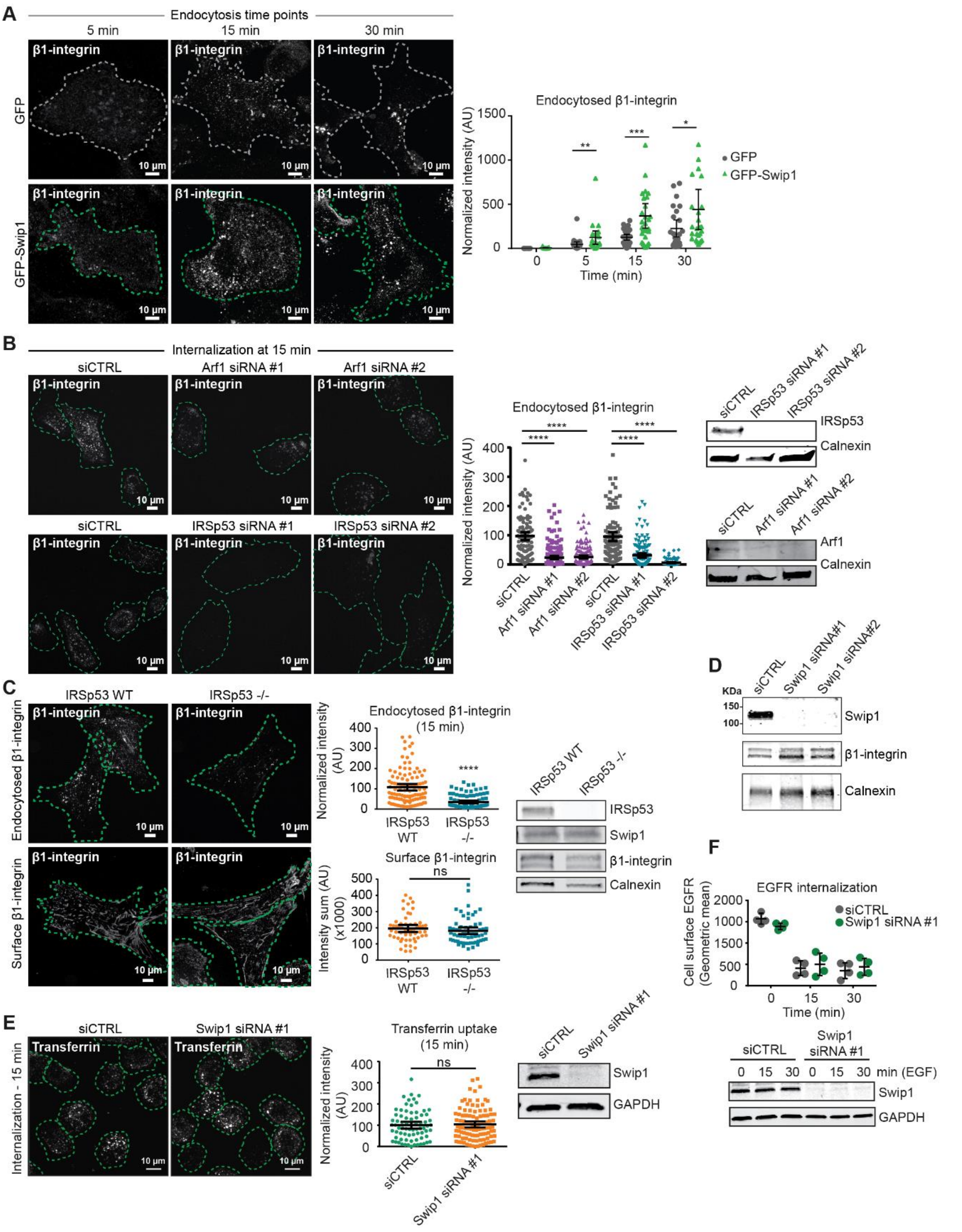


**Extended Data Fig. 5. Swip1 regulates β1-integrin endocytosis via the CG pathway.**
**(A)** Ectopic expression of GFP-swip1 increases β1-integrin endocytosis. Representative micrographs of β1-integrin uptake in GFP and GFP-swip1 expressing cells and quantification of integrin uptake at 5, 15 and 30 min. **(B)** Representative micrographs and quantification of β1-integrin uptake at the 15 min time point in control- or IRSp53- or Arf1-silenced MDA-MB-231 cells. Representative immunoblots of cell lysates blotted as indicated, calnexin is included as a loading control. **(C)** Representative micrographs and quantification of endocytosed (15 min uptake) or surface (no endocytosis or acid wash) of murine β1-integrin (9EG7 antibody) in isogenic IRSp53−/− MEFs: IRSp53-KO-pBABE (-/-) and IRSp53-KO-pBABE-IRSp53 (WT) in which the expression of IRSp53 has been restored. Representative immunoblots of cell lysates blotted as indicated, calnexin is included as a loading control. **(D)** Representative immunoblots of control- and swip1-silenced MDA-MB-231 cell lysates blotted as indicated, calnexin is included as a loading control. **(E)** Representative micrographs and quantification of transferrin uptake at the 15 min time point in control or swip1 silenced MDA-

MB-231 cells. Representative immunoblots of cell lysates blotted as indicated, calnexin is included as a loading control. **(F)** Cell-surface labelled EGFR uptake in control- and swip1-silenced MDA-MB-231 cells treated with 10 ng/ml EGF was analyzed using flow cytometry in non-permeabilized cells (mean ± SD, 4-5 measurements from 2 experiments).

Number of analyzed cells: GFP, 10, 22, 30 and 24 and GFP-Swip1, 15, 22, 32 and 26, respectively. Plots are mean ± 95 % CI, $n = 3$ experiments, *$P < 0.05$; **$P < 0.005$, ***$P < 0.0005$, Mann-Whitney test.

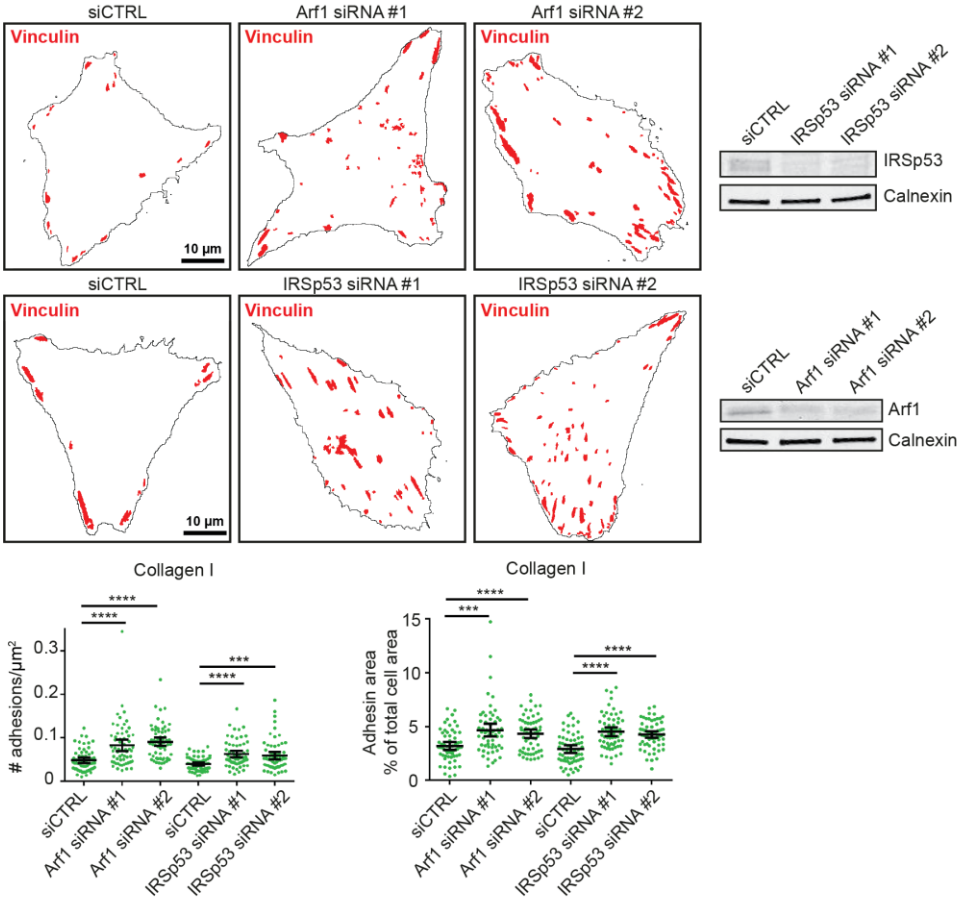


**Extended Data Fig. 6. Arf1 and IRSp53 regulate focal adhesion formation.**
Representative masks of vinculin-containing focal adhesions in control-, IRSp53- or Arf1-silenced MDA-MB-231 cells and quantification of adhesion number and total area per cell in cells plated on collagen I. Data are mean ± 95 % CI, *** $P < 0.0005$, $n = 3$ experiments. Representative immunoblots to validate IRSp53 and Arf1 silencing are shown.

**EFHD2 mRNA level**

| | | < median | | > median | | total samples | p |
|---|---|---|---|---|---|---|---|
| | | # samples | % samples | # samples | % samples | | |
| molecular subtype | HER2+ | 10 | 34.5% | 19 | 65.5% | 29 | 0.002 |
| | luminal | 72 | 58.5% | 51 | 41.5% | 123 | |
| | TNBC | 11 | 29.7% | 26 | 70.3% | 37 | |
| total | | | | | | 189 | |
| grading | 1 | 12 | 60.0% | 8 | 40.0% | 20 | 0.049 |
| | 2 | 47 | 58.8% | 33 | 41.3% | 80 | |
| | 3 | 38 | 41.3% | 54 | 58.7% | 92 | |
| total | | | | | | 192 | |

**Extended Data Fig. 7. Swip1 mRNA expression correlates with a more metastatic breast cancer phenotype.**
Analysis of swip1 mRNA expression using qPCR in 192 breast tumors. Tumors were categorized into two groups: high swip1 expression (higher than the median of the entire collection) or low swip1 expression (lower expression than the median). The $\chi 2$-test was used for statistical analyses. Swip1 mRNA levels based on tumor type and tumor grade are shown.

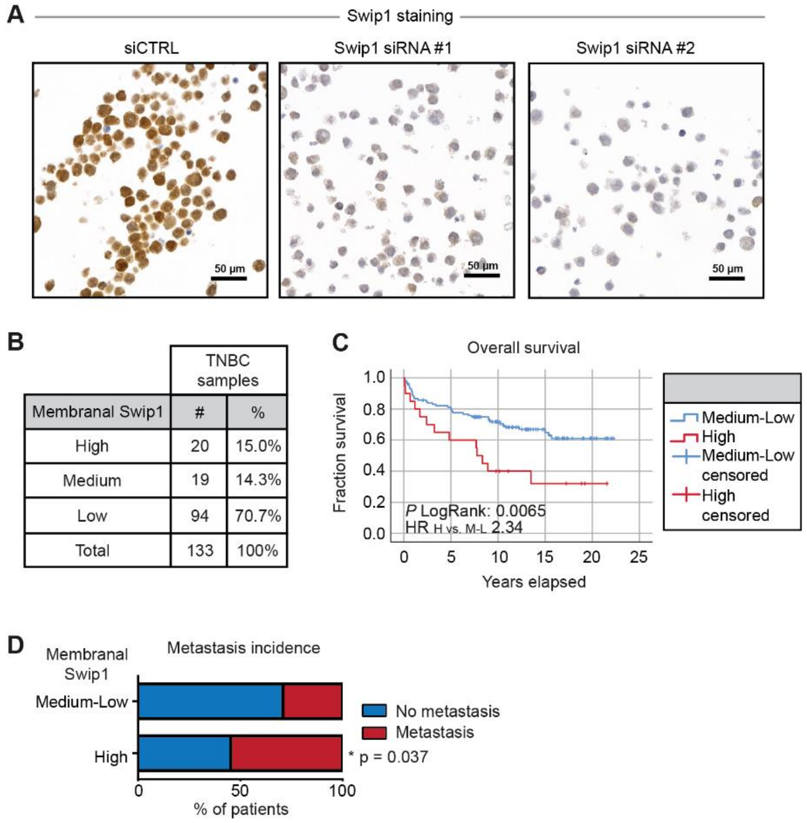


**Extended Data Fig. 8. Swip1 immunostaining at the plasma membrane correlates with higher death risk and metastasis incidence in TNBC.**
**(A)** Validation of anti-swiprosin-1 antibody specificity in IHC was carried out with agarose embedded and *Formalin*-Fixed Paraffin-Embedded cell pellets from control- or swip1-silenced MDA-MB-231 cells (siRNA #1 and #2). **(B)** Quantification of the percentage of breast cancer tumours from TNBC tissue microarray with high, medium or low swiprosin-1 staining at the plasma membrane. **(C)** Kaplan-Meier plot shows overall survival of 133 triple negative breast cancer patients with high (H, red) or medium-low (M-L, blue) membranal staining of swip1 in their tumor center sample. The hazard ratio of high vs. medium-low swiprosin-1 immunopositivity was 2.34 (95% CI 1.25 to 4.41). The hazard ratios after adjustment for Ki67, 2.51 (95% CI 1.26 to 5.01); for tumor size, 2.12 (95% CI 1.08 to 4.18); for lymph node metastasis status 2.26 (95% CI 1.19 to 4.31) and for tumor grade 2.44 (95% CI 1.25 to 4.74). **(D)** Lymph node metastasis incidence in 130 TNBC patients from the same cohort, Fisher's exact test p=0.037. A higher proportion of patients with high membranal staining of swiprosin-1 (>=80%) had lymph node metastasis.

**Supplementary Table 1 (provided as a separate file).** Log10 transformed ratios of the proteins identified by mass spectrometry.

**Supplementary Table 2.** Antibodies used in this study.

| **Antibody** | **Manufacturer** | **Catalogue nr** | **Application** | **Dilution** |
|---|---|---|---|---|
| Swiprosin-1 (EFHD2) | Everest Biotech | EB06552 | WB<br>PLA | 1:1000<br>1:100 |
| Swiprosin-1 (EFHD2) | Atlas Antibodies | HPA048961 | WB<br>IHC | 1:1000<br>1:200 |
| Rab21 | Self made [22] | - | WB<br>IF | 1:500<br>1:50 |
| Integrin β1 P5D2 | Developmental Studies Hybridoma Bank | - | WB<br>IF | 1:1000<br>1:100 |
| Integrin β1 | BD Biosciences | 610468 | WB | 1:1000 |
| IRSp53 (BAIAP2) | Atlas Antibodies | HPA023310 | WB | 1:1000 |
| Arf1 | GeneTex | GTX113385 | WB | 1:1000 |
| GFP | Invitrogen | A11122 | WB | 1:1000 |
| GFP | Abcam | Ab1218 | IF | 1:100 |
| Calnexin | Enzo Life Sciences | ADI-SPA-865-F | WB | 1:1000 |
| GAPDH | HyTest | 5G4MaB6C5 | WB | 1:2000 |
| β-actin | Sigma | A1978 | WB | 1:1000 |
| Integrin β1 12G10 | Supernatant from hybridoma provided by K.M. Yamada (NIH, Bethesda, MD) | - | IF | 1:10 |
| Integrin β1 9EG7 | BD Biosciences | 553715 | IF | 1:100 |
| Clathrin | Abcam | Ab21679 | IF | 1:100 |
| AP2 | Thermo Scientific | MA1-064 | IF | 1:100 |
| Caveolin | Santa Cruz | SC-894 | IF | 1:100 |
| MHCI | Sigma | SAB4700637 | IF | 1:200 |
| Vinculin | Sigma | V9131 | IF | 1:100 |
| Rab5 | Cell Signaling Technology | 3547S | IF | 1:100 |
| Rab7 | Cell Signaling Technology | 9367S | IF | 1:100 |
| Rab11 | Cell Signaling Technology | 5589S | IF | 1:100 |
| EGFR | Upstate | 05-101 | FACS | 1:100 |
| HA-tag | Cell Signaling Technology | 3724S | IF | 1:100 |

**Supplementary Video 1.**
MDA-MB-231 cells transiently co-expressing V1-Rab21 and V2-swip1 (A) or expressing Venus control (B) were imaged using TIRF microscopy (84.5° angle) every 500 ms for 5 min.

**Supplementary Video 2.**
MDA-MB-231 cell transiently co-expressing V1-Rab21, V2-swip1 and mcherry-IRSp53, imaged using TIRF microscopy (84.5° angle) every 500 ms for 2.5 min. The white squares show regions of interest (ROI) where V1-Rab21+V2-swip1 puncta (green) travel to reach mcherry-IRSp53 puncta (magenta) before exiting the TIRF plane. One of the ROI is depicted as a timelapse in Figure 1M.

**Supplementary Video 3.**
MDA-MB-231 cell transiently expressing GFP-swip1 and treated with SiR-Actin, imaged using semi-superresolution microscopy.

**Supplementary Video 4.**
MDA-MB-231 cell (siCTRL) stably expressing GFP-Rab21 and imaged using TIRF microscopy (76.43° angle) every 500 ms for 2 min.

**Supplementary Video 5.**
MDA-MB-231 cell (Swip siRNA #2) stably expressing GFP-Rab21 and imaged using TIRF microscopy (76.43° angle) every 500 ms for 2 min.
